# Large scale CRISPR genome engineering reveal principles of single nucleotide microhomology in precision editing

**DOI:** 10.64898/2026.11.16.766972

**Authors:** Xin Shi, Chenyu Li, Jingwei Li, Yuzhe Liu, Qiang Wu

**Author notes:** These authors contributed equally (Xin Shi, Chenyu Li).

## Abstract

The CRISPR/Cas9 system has emerged as a revolutionary tool with diverse applications in genome engineering; however, the role of nanoscale hydrogen bonds between single nucleotides in precision editing remains unknown. Here we systematically compared A-rich and G-rich PAM-recognizing Cas9 editors through comprehensive analyses of ∼37,000 targeting sites with about half million libraries of ∼0.36 billion types of editing outcomes. We showed that the A-rich Cas9 editors exhibit stronger editing outcome predictability in comparison with the conventional Cas9 editor of the G-rich PAM. In addition, the A-rich Cas9 editors induce a higher frequency of staggered cleavages, resulting in predictable editing with donor-independent templated insertions. Moreover, we identified distinct DNA sequence motifs for predictable insertions with the A-rich Cas9 editors. In particular, we revealed single nucleotides at the −4 position as MMEJ microhomologies in precision engineering via two or three hydrogen bonds with A-rich or G-rich PAM, respectively. Furthermore, we found that PolQ plays a key role in enhanced random insertions. Finally, we developed a discrete-space diffusion model CRIfuser of deep learning for predicting editing outcomes with staggered cleavage. Together, these findings establish PAM-dependent cleavage geometry as a tunable determinant of Cas9 editing outcomes and provide a framework for predictive and precision genome engineering.

## Introduction

The rapid advancement of gene targeting or genome editing technologies has greatly accelerated reverse genetics, which serves as a key method for uncovering gene functions and biological mechanisms[1,2]. In particular, the CRISPR/Cas9 system (clustered regularly interspaced short palindromic repeat, CRISPR), which originates from the adaptive immune system of bacteria and archaea, has been widely used in genome editing owing to its high efficiency and low cost[2]. Specifically, SpyCas9 programmed by a synthetic single-guide RNA (sgRNA) generates blunt and staggered double strand break (DSB) ends at cleavage sites[2–6]. These DSB ends are subsequently repaired by cellular machineries, often resulting in insertions and deletions (indels)[7]. For example, ligation of two DSB ends filled in by Polλ of 5’ overhangs, which are generated by the staggered endonuclease cleavage activity of SpyCas9, results in predictable donor-independent templated insertions[5,6,8–12].

The widely-used SpyCas9 requires a G-rich NGG protospacer adjacent motif (PAM) for target recognition[13]. The PAM sequence, a short motif downstream of the targeting site, is recognized by the PAM-interacting (PI) domain which triggers the crucial DNA duplex unwinding and R-loop formation prior to targeted cleavage[14,15]. Specifically, the PI domain of SpyCas9 utilizes residues R1333 and R1335 to recognize the two guanines (G) in the major groove of the NGG PAM[15]. However, simply mutating these two arginine residues into glutamine (Q), which typically recognizes adenine (A), fails to switch PAM from NGG to NAA, suggesting that PAM recognition is not solely the major groove interactions with the two Cas9 residues[16]. To this end, a PI-domain-engineered variant was developed by introducing seven amino-acid mutations for recognition of the A-rich NAA PAM[17]. However, due to its low editing efficiency, this variant has not been widely used.

Here we conducted large-scale high-throughput analyses to characterize the properties of A-rich PAM-recognizing Cas9 editors in precision editing. In addition, we uncovered a role of PolQ in insertional editing through massively parallel profiling. Finally, by a diffusion-based machine learning approach, we developed the CRIfuser AI program to predict the editing outcomes for any targeting site and revealed the crucial PAM-flanking nucleotides for Cas9 staggered cleavage.

## Materials and methods

### Oligonucleotide design for the NAA PAM sgRNA library with the NGG control library

To systematically analyze the properties of Cas9 editors that recognize A-rich and G-rich PAMs, two oligonucleotide pools were designed and synthesized on an array platform by GenScript. Each oligonucleotide was designed to sequentially include the following components: (i) a 19-nt primer binding site, (ii) an sgRNA sequence, (iii) a *Bsm*BI restriction enzyme recognition site, (iv) a 20-nt barcode sequence (Barcode 1), (v) a second *Bsm*BI site, (vi) a 44-nt target sequence containing either the NAA or NGG PAM (with the cleavage site positioned at the center of the target sequence), (vii) an 18-nt barcode sequence (Barcode 2), and (viii) a 20-nt primer binding site.

Barcode 1, located between the two *Bsm*BI sites, served to reduce template switching during oligonucleotide amplification. Barcode 2, located downstream of the target site, was designed to identify its corresponding sgRNA sequence, enabling the precise demultiplexing of high-throughput sequencing reads even if the target sequences were truncated after editing. All barcode sequences were generated using DNA Barcodes (https://github.com/feldman4/dna-barcodes)[18], ensuring a Hamming distance of 2 (at least two mismatches) between any two barcodes.

Oligonucleotides were filtered out if they contained internal *Bsm*BI restriction sites, primer sequences within the sgRNA or target regions, or homopolymeric tracts of five consecutive bases (AAAAA, TTTTT, CCCCC, GGGGG). Finally, an extra G was added immediately upstream of the sgRNA sequence to promote robust expression driven by the U6 promoter.

Each library comprised 37,266 sgRNA-target pairs that differed only in their PAM sequences: one contained an NAA PAM (library A) and the other an NGG PAM (library G). Owing to synthesis capacity constraints, each library was subdivided into three sub-libraries. The nucleotide composition of the synthesized sequences was well-balanced, featuring an average GC content of approximately 50%. The sequences of all synthesized oligonucleotides are listed in Supplementary Table S1.

### Construction and quality control of sgRNA lentiviral libraries

sgRNA plasmid libraries were generated using a two-step cloning strategy. First, the synthesized oligonucleotide pools were amplified using the minimum number of PCR cycles determined by qPCR and cloned by Gibson assembly into the linearized lentiCRISPR backbone. This backbone was derived from lentiCRISPR (Addgene #52961) by deleting the Cas9-coding sequence and the sgRNA scaffold sequence. The assembled products were transformed into Endura electroporation-competent cells, and colonies were pooled after overnight growth at 32°C. Plasmid libraries were extracted using the QIAGEN Plasmid Maxi Kit (Qiagen, 12165). Sanger sequencing of randomly selected colonies confirmed successful library construction, with an initial library representation exceeding 100-fold.

Second, the sgRNA scaffold was inserted into the initial libraries. The initial plasmid libraries were linearized with *Bsm*BI (NEB, E1602S), dephosphorylated, gel-purified and ligated with *Bsm*BI-digested scaffold amplicons. The final ligation products were transformed into competent cells and recovered as described above. Final libraries were validated by Sanger sequencing and maintained more than 50-fold representation. All primers used for sgRNA plasmid library construction and scaffold amplification are listed in Supplementary Table S2.

Library quality was assessed by high-throughput sequencing with a first-round low-cycle PCR for the synthesized oligonucleotide region, followed by a second-round PCR to incorporate Illumina-compatible P5 and P7 adapters. After purification, amplicons were quantified using Qubit fluorescent dye (Vazyme EQ121-01), and sequenced on the Illumina NovaSeq 6000 platform at more than 5,000-fold coverage. All primers used for library quality assessment are listed in Supplementary Table S2.

### Cell culture

HEK293T cells (human embryonic kidney cells, female, ATCC CRL-3216) and all HEK293T-derived cell lines generated in this study were cultured in Dulbecco’s Modified Eagle’s Medium (DMEM; Gibco, #C11995500BT) supplemented with 10% fetal bovine serum (FBS; Gibco, #260100741) and 1% penicillin/streptomycin (Gibco, #15140122). Cells were maintained at 37 °C in a humidified incubator with 5% CO_2_.

### Lentivirus production for sgRNA libraries

sgRNA plasmid libraries were packaged in HEK293T cells as previously described[19–21]. Briefly, when cells reached ∼60% confluence, they were co-transfected with the transfer plasmid (sgRNA plasmid libraries) and packaging helper plasmids (psPAX2 and pMD2.G) at a 4:3:2 mass ratio using Lipofectamine 3000 (Invitrogen, L3000015). The medium was replaced with fresh growth medium supplemented with 30% FBS at 18h post transfection. Lentivirus-containing supernatants were collected at 40, 50, and 60h after transfection. The supernatants were then filtered through 0.45-µm low-protein-binding membranes and subsequently concentrated by ultracentrifugation at 12,000g for 1.5h at 4 °C. The viral pellets were resuspended in PBS and stored at −80 °C until use. Viral titers were determined using a live-cell infection assay. Briefly, after infecting cells with serial dilutions of virus in the presence of 8 µg/ml polybrene followed by selection with 2 µg/ml puromycin, viral titers were calculated from the number of antibiotic-resistant cells.

### Optimization of sgRNA scaffold for A-rich PAM-recognizing SpyMac and iSpyMac Cas9 editors

To improve the editing efficiency for A-rich PAM-recognizing Cas9 editors and determine the sgRNA scaffold used for pooled profiling, we compared conventional, optimized, and Smac (*Streptococcus macacae*) sgRNA scaffolds. Optimized and Smac scaffold-containing vectors were generated by replacing the conventional scaffold in the sgRNA expression backbone. Briefly, the Cas9-deleted backbone derived from lentiCRISPR (Addgene #52961) was linearized with *Pac*I (NEB, R0547V) and *Pme*I (NEB, R0560V). Scaffold-containing fragments were amplified by overlapping PCR and the final sgRNA-expression plasmids were assembled by Gibson assembly. All plasmids were confirmed by Sanger sequencing.

The efficiency of optimized and Smac scaffolds were evaluated at ten human genomic loci for SpyMac and iSpyMac Cas9 editors with the conventional sgRNA scaffold as a control. To generate constructs for each sgRNA transcription, a pair of two complementary oligonucleotides were annealed to form double-stranded DNA (dsDNA) with 5’ overhangs of ‘CACC’ and ‘AAAC’ and the annealed DNA was inserted into *Bsm*BI-linearized sgRNA plasmid backbones. The sgRNA-expressing plasmids were co-transfected with either SpyMac or iSpyMac expression plasmids into HEK293T cells. Two days after transfection, cells were selected with 2 µg/ml puromycin for additional four days. Genomic DNA was extracted with Wizard Genomic DNA Purification Kit (Promega A1120) following the manufacturer’s instructions. Briefly, after collecting the cell pellet, the cells were resuspended with Nuclei Lysis Solution and lysed at 37°C for 30 min. Then, the proteins were precipitated with Protein Precipitation Solution and removed by centrifugation at 12,000 rpm for 10 min. The supernatant containing the genomic DNA was then extracted with isopropanol followed by washing with 70% ethanol. The DNA pellet was finally dissolved in nuclease-free water. After PCR amplification of the ten targeting loci, editing efficiencies were analyzed by Sanger sequencing using ICE2 (https://github.com/synthego-open/ice)[22]. All primer sequences used for sgRNA scaffold optimization are listed in Supplementary Table S2.

### Construction of Cas9 plasmids and lentivirus production

To improve lentiviral titers, Cas9 and sgRNA were cloned in separate vectors. SpyCas9, SpyMac, and iSpyMac plasmids were constructed from lenti dCAS-VP64 backbone (Addgene #61425) using Gibson assembly. For SpyCas9, the backbone was linearized with *Not*I and *Xho*I, and the full-length SpyCas9 coding sequence was assembled with the vector fragments. The SpyMac plasmid was generated using the same backbone by replacing the SpyCas9 PAM-interacting domain with a synthesized Smac-derived PI-domain fragment. For iSpyMac plasmid construction, point mutations of R221K and N394K were introduced into SpyMac plasmid by overlapping PCR. Finally, the SpyCas9, SpyMac, and iSpyMac plasmids were packaged into lentiviral particles with the two helpers using the same lentivirus production workflow as described above. All final constructs were validated by Sanger sequencing, and the primer sequences are listed in Supplementary Table S2.

### Generation and validation of HEK293T-SpyCas9, HEK293T-SpyMac, HEK293T-iSpyMac, HEK293T-ΔPolQ, HEK293T-ΔPol, and HEK293T-ΔHel stable cell lines

To establish stable Cas9-expressing single cell clones, HEK293T cells were transduced with lentiviruses encoding SpyCas9, SpyMac, or iSpyMac at an MOI of 4 in the presence of 8 µg/ml polybrene. After 24h, cells were selected with 4 µg/ml blasticidin (Invitrogen) for 7-10 days. Blasticidin-resistant cells were subjected to limiting dilution in 96-well plates to obtain monoclonal cells. We screened 671, 596, and 548 single-cell clones for SpyCas9-, SpyMac-, and iSpyMac-expressing stable cell clones, respectively. The primers used for genotyping are listed in Supplementary Table S2. For each Cas9 editor, protein expression levels were evaluated by western blotting as described below. Two clones with high and comparable Cas9 expression were selected for sgRNA library transduction.

PolQ-deficient HEK293T lines were derived from SpyCas9-expressing cell line. To generate HEK293T-ΔPolQ single-cell clones, dual sgRNAs were employed to induce DNA fragment deletion spanning the *PolQ* exons. In contrast, HEK293T-ΔPol and HEK293T-ΔHel single-cell clones were established via co-transfection of sgRNA-expressing plasmids and donor templates designed to delete specific domain-coding sequences. Following transfection, cells were selected with 2 µg/ml puromycin, and monoclonal populations were isolated through limiting dilution. The genotypes of all candidate clones were validated by PCR followed by Sanger sequencing. All primer sequences utilized for construction and screening are provided in Supplementary Table S2. From the initial screening of 356, 306, and 319 single-cell clones, we identified 91, 47, and 36 heterozygous clones for HEK293T-ΔPolQ, HEK293T-ΔPol, and HEK293T-ΔHel, respectively. Ultimately, two independent homozygous clones were successfully isolated and confirmed for each condition.

### sgRNA library transduction and genomic DNA extraction

Stable Cas9-expressing and PolQ-deficient single-cell clones were transduced with the corresponding sgRNA lentiviral libraries at an MOI of 0.6 in the presence of 8 µg/ml polybrene to ensure single-copy integration. Two independent biological replicates were performed for each stable cell line, with more than 1,000-fold library coverage. At 24h post-transduction, successful transducing cells were selected with 2 µg/ml puromycin for 7-10 days. After selection, cells were collected in numbers sufficient to maintain at least 1,000-fold library representation. The genomic DNA was extracted using the Wizard Genomic DNA Purification Kit (Promega, A1120) following the manufacturer’s instructions. To obtain highly purified genomic DNA, the extracted DNA was passed through a miniprep column.

### Deep sequencing of repair outcomes

To evaluate DNA repair outcomes in a high-throughput manner, a two-step PCR amplification strategy was adapted from established protocols for sgRNA library qualification mentioned above. Following genomic DNA extraction and quantification, the total number of PCR reactions was determined by the maximum template input (25 µg per 50 µl reaction). To ensure library representations of at least 1,000-fold coverage, the number of parallel reactions was scaled accordingly. The first-round PCR round was conducted to capture indel profiles using the following conditions: initial denaturation at 98 °C for 30s, followed by 20-24 cycles of 98 °C for 10s, 60 °C for 20s, and 72 °C for 30s, and a final extension at 72 °C for 5 min. Amplicons were subsequently pooled, and 5 µl of the pooled PCR product was utilized as templates for the second-round amplification. The second-round PCR was performed using the following conditions: 98 °C for 30s, 8 cycles of 98 °C for 10s, 60 °C for 20s, and 72 °C for 30s, followed by a final extension at 72 °C for 3 min. The resulting libraries were purified using the QIAquick PCR Purification Kit (Qiagen, 28104), and quantified using Qubit fluorescent dye (Vazyme, EQ121-01). All libraries were sequenced on the Illumina NovaSeq 6000 platform, maintaining a minimum sequencing depth of 1,000-fold coverage per target site. All primer sequences are provided in Supplementary Table S2.

### Western blot

The expression levels of Cas9 editors were validated via western blotting. Briefly, upon reaching ∼80% confluence in 6-well plates, cells were harvested by detachment with 0.25% trypsin-EDTA (Gibco, 25200056). Following the removal of the supernatant, the cell pellets were resuspended in pre-cooled RIPA lysis buffer (Sangon, C510006) and incubated on ice for 15 min. Cell lysates were subsequently centrifuged at 12,000 rpm for 10 min at 4 °C. The resulting supernatants were collected and stored at −80 °C for subsequent analysis.

Protein concentrations were determined using a BCA protein assay kit (Beyotime, P0011) according to the manufacturer’s instructions. For each sample, 20 µg of total protein was mixed with 1×SDS loading buffer, loaded onto 4-12% Bis-Tris gels, and separated in 1×MOPS SDS running buffer (Sangon, C691402-0001) at 160 V for 45 min. Following electrophoresis, proteins were transferred onto nitrocellulose membranes at 200 V for 150 min. The membranes were then blocked with 5% non-fat milk in 1× PBST for 1h at room temperature. After three washes in 1× PBST, the membranes were incubated overnight at 4 °C with the primary antibody specific to the N-terminus of SpyCas9 (BioLegend, 844301). Owing to high sequence similarity of N-terminus among SpyCas9, SpyMac, and iSpyMac, the antibody can be used to detect all three editors. The membranes were subsequently washed three times in 1× PBST and incubated with the secondary antibody at 4 °C for 16h. After the removal of unbound antibodies via three additional 1× PBST washes, protein bands were visualized and quantified using the Odyssey Imaging System (LI-COR Biosciences).

### High-throughput sequencing data analyses

We developed and utilized NAA PAM (https://pypi.org/project/naapam/), a custom bioinformatic pipeline, to accurately quantify CRISPR/Cas9-mediated editing outcomes (Supplementary Note S3). A critical challenge in such analysis is that oligonucleotide synthesis errors can confound the estimation of true editing profiles. For instance, a 1-bp deletion at −5 position resulting from a synthesis error is often indistinguishable from a 2-bp PAM-distal deletion accompanied by a 1-bp templated insertion. Because templated insertions induced by staggered cleavage are relatively infrequent, even minimal background synthesis error can lead to a substantial overestimation of templated insertion frequencies.

To address this, we implemented a robust filtering strategy. First, we utilized sequencing reads from control samples (without Cas9) as reference templates for mapping to eliminate discrepancies between designed and synthesized sequences. Second, we applied a statistical model to exclude targeting sequences with aberrant high PAM-distal deletion frequencies for any given templated insertion size. The expected relative frequencies of PAM-distal deletions, relative to blunt-end junctions, was estimated by fitting a linear model of the log-transformed relative abundances against deletion sizes. Targeting sequences with the relative PAM-distal deletion frequencies exceeding the upper bound of the 95% prediction interval were flagged as outliers and excluded from subsequent analyses.

Reads were filtered based on the structural integrity of essential elements, including primers, spacers, scaffolds, barcodes, protospacers, and PAM sequences. Furthermore, only reads with matching between spacer and protospacer in the control cells with no editor were retained, ensuring that the analyzed sequences represented valid biological templates available for Cas9 editing.

Following preprocessing, reads were aligned to their references using a tripartite alignment strategy with a few modifications[11]. Each read was partitioned into three segments: upstream, middle, and downstream. The upstream and downstream segments were mapped independently to the reference; notably, the 5’ end of the downstream segment was permitted to map to the nucleotides upstream of the cleavage site to capture the templated insertions resulting from staggered cleavage. The middle segment was treated as an unmapped random insertion. This partition and mapping of the upstream and downstream segments were integrated into an optimized chimeric alignment engine powered by dynamic programming.

To further distinguish Cas9-induced edits from the background noise, such as PCR artifacts or residual synthesis errors, the final indel frequency was calculated by subtracting the background indel frequency observed in the control library using the following formula:

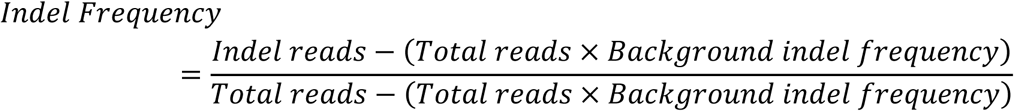

As a final quality-control threshold, targeting sequences were excluded if they exhibited an adjusted indel frequency below 8% or a total read depth of fewer than 100 reads.

### Repair profile analysis

Sequencing reads were binned for each sgRNA-target pair based on unique 18-bp barcode sequences. Following deduplication and demultiplexing, the reads were aligned to their respective reference sequences to characterize repair profiles. We categorized indel outcomes into eight distinct groups: wild type (WT), templated insertions (TI) only, random insertions (RI) only, deletions only, and all possible combinations thereof (TI + deletion, TI + RI, RI + deletion, and TI + RI + deletion).

Deletions were further classified as either microhomology-mediated end joining (MMEJ) or non-MMEJ deletions. Non-MMEJ deletions were subdivided based on their position relative to the cleavage site: PAM-distal deletions, PAM-proximal deletions, or bidirectional deletions (spanning the cleavage site). Furthermore, repair outcomes were characterized by size, specifically focusing on insertions of 1-10 bp and deletions of 1-30 bp. Deletions were also mapped by their precise coordinates (start and stop sites). To facilitate comparisons between different Cas9 variants, unless otherwise specified, all indel counts were normalized against the total number of mutated reads for each sgRNA-target pair.

To quantify the diversity of DNA repair outcomes, we used the Shannon entropy index, calculated as follows:

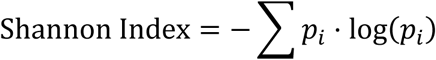

where *p_i_* represents the frequency of a specific indel event *i* for a given target sequence.

DNA motifs associated with editing efficiency, templated insertions, and MMEJ deletions were analyzed using kpLogo. The analysis was performed with default parameters using the k-mer length of 1 as previously described[23]. We examined a predefined 20-nucleotide (nt) window immediately upstream of the PAM sequence. To ensure the motifs accurately reflected the relative abundance of specific repair outcomes, input sequences were weighted according to their observed frequencies of editing efficiency, templated insertions, or MMEJ deletions.

### Microhomology definition

To accurately characterize MMEJ-mediated repair, CRISPR/Cas9 editing outcomes were aligned to the target site sequence chimerically. Each chimeric alignment was decomposed of two contiguous blocks and an optional random insertion between them. We defined microhomology (MH) as the longest common subsequence shared between the context at the 3’ terminus of the upstream block and the 5’ terminus of the downstream block. When multiple optimal alignments were generated, we systematically evaluated all candidates and prioritized the one yielding the maximum microhomology length. This approach ensures that MMEJ-driven repair signatures are not underestimated due to arbitrary alignment scoring, thereby providing a more biologically relevant representation of the DNA repair landscape.

### AI model for predicting editing profiles

To predict the editing outcomes for SpyMac and iSpyMac, we developed CRIfuser, a discrete-space diffusion[24] probabilistic model designed to characterize the joint 2D distribution of PAM-distal and PAM-proximal DNA ligation ends. CRIfuser treats repair outcomes as a discrete categorical distribution across a defined 2D coordinate space (Supplementary Note S1). The core architecture employs a U-Net backbone integrated with a linear denoising scheduler 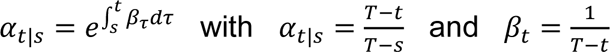 for the reverse diffusion process. Rather than regressing the final profile into a single pass, the model iteratively refines a randomly initialized outcome (*x_T_*) into a final prediction (*x*_0_). At each time step *t*, the U-Net predicts the transition probability *p*(*x_t_*_−1_|*x_t_*, *C*) conditioned on input features (*C*). This iterative refinement allows the model to capture complex, non-linear dependencies between the two ligation ends that are often missed by single-pass architectures.

The model functions within a 32×32 grid representing the 2D diffusion space, with a PAM-distal range of −25 bp to +6 bp and a PAM-proximal range from −6 bp to +25 bp relative to the cleavage site. This configuration enables the prediction of deletions up to 25 bp and templated insertions up to 6 bp. The input to the U-Net architecture is an 11-channel tensor (11×32×32) that integrates multi-layered biological and positional information: four channels each for one-hot encoded PAM-distal and PAM-proximal sequences (totally eight channels, derived from a 50bp window surrounding the cleavage site), a single channel for a 25×25 microhomology matrix (zero-padded to 32×32) to facilitate the modeling of MMEJ-mediated repair, a one-hot 2D tensor marking the precise cleavage location, and a channel representing the editing state from the preceding diffusion step. Within the U-Net backbone, “down” modules incorporate sinusoidal embeddings to encode time-step information, guiding the denoiser through the iterative stages of reverse diffusion.

CRIfuser was trained on large-scale editing datasets from SpyCas9, SpyMac, and iSpyMac, and benchmarked against both baseline architectures including RoFormer, LSTM, CNN, MLP, XGBoost, and ridge regression and previous models including inDelphi[25], FOREcasT[26], and Lindel[27]. The optimization objective combined local and global loss components to ensure robust predictive performance. Specifically, we utilized an Evidence Lower Bound (ELBO) based on importance sampling:

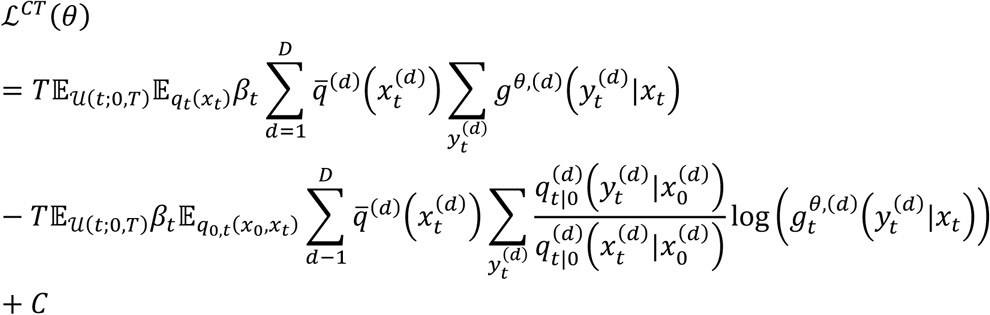

as the local loss to optimize reverse diffusion transitions, complemented by a conditional cross-entropy global loss:

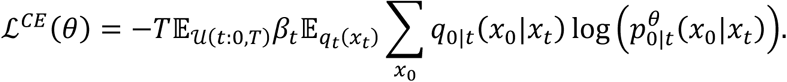

To enhance training stability and accelerate convergence, a time step zero reparameterization was implemented; rather than calculating loss based on a single sample, the model predicts the denoised profile directly from the noisy state at each step, facilitating a more consistent gradient flow. To handle the resulting high-dimensional tensor operations, we optimized the tensor contraction order, ensuring feasible computation times. All models were trained using the hyperparameters in Supplementary Table S3 for 33 epochs, incorporating a linear warmup phase during the first 3 epochs. Final model selection was guided by the negative log-likelihood (NLL) and mean squared error (MSE) relative to the ground-truth editing profiles derived from deep sequencing reads.

Since diffusion models are inherently stochastic samplers, generating a stable population-level editing profile through individual outcome sampling is computationally inefficient and prone to sampling noise. To solve this problem, instead of sampling a single path, we computed the transition across all possible outcomes at each step, weighted by their marginal probabilities. For interpretability analysis (e.g., SHAP values), where thousands of forward passes are required, we further implemented an interpolation-based grid sampling method. By sampling a subset of the 32×32 space and distributing weights via distance-based linear interpolation, we maintained high predictive accuracy while drastically reducing the computational overhead.

## Results

### A-rich PAM-recognizing Cas9 editors exhibit higher precise and predictable insertions

Two Cas9 editors of SpyMac and iSpyMac derived from *Streptococcus macacae* (SmacCas9) was shown to recognize A-rich PAM sequences[28]. We first optimized the sgRNA scaffold to improve editing efficiencies (Supplementary Fig. S1)[22]. We then constructed a high-throughput CRISPR library containing 37,266 targeting sites each with an NAA PAM; for comparative analyses, we also constructed a similar library containing 37,266 identical targeting sites but each with an NGG PAM as controls (Supplementary Fig. S2). We found that the CRISPR libraries are robust and efficient (Supplementary Fig. S3).

We then comprehensively analyzed insertion profiles upon cleavage with A-rich and G-rich PAM-recognizing Cas9 editors. Insertional editing outcomes can be categorized into templated insertions and non-templated random insertions (Supplementary Fig. S3C). We characterized the cleavage profiles of 37,266 targeting sites and found that the A-rich PAM-recognizing Cas9 editors of both SpyMac and iSpyMac exhibited significantly higher percentage of 1-10bp templated insertions than that of the G-rich SpyCas9 (Fig. 1A). This suggests that the A-rich PAM-recognizing editors preferentially generate more overhangs via staggered cleavage. Moreover, both the A-rich and G-rich PAM-recognizing Cas9 editors generate remarkable overhangs of 4-10 bp (Fig. 1A), in addition to 1-3 bp overhangs previously known[5,6,8]. For example, the percentages of templated 4-bp insertions in total 4-bp insertion reads (31.04%, 53.01%, 46.64% for SpyCas9, SpyMac, and iSpyMac, respectively) are far more than what would be expected from random insertions (Fig. 1A). Finally, we found that the A-rich and G-rich Cas9 editors have distinct editing patterns (Supplementary Fig. S4). However, the frequencies of templated insertions in total edited reads drop dramatically for 5-10bp (Supplementary Fig. S5A).

**Figure 1.**
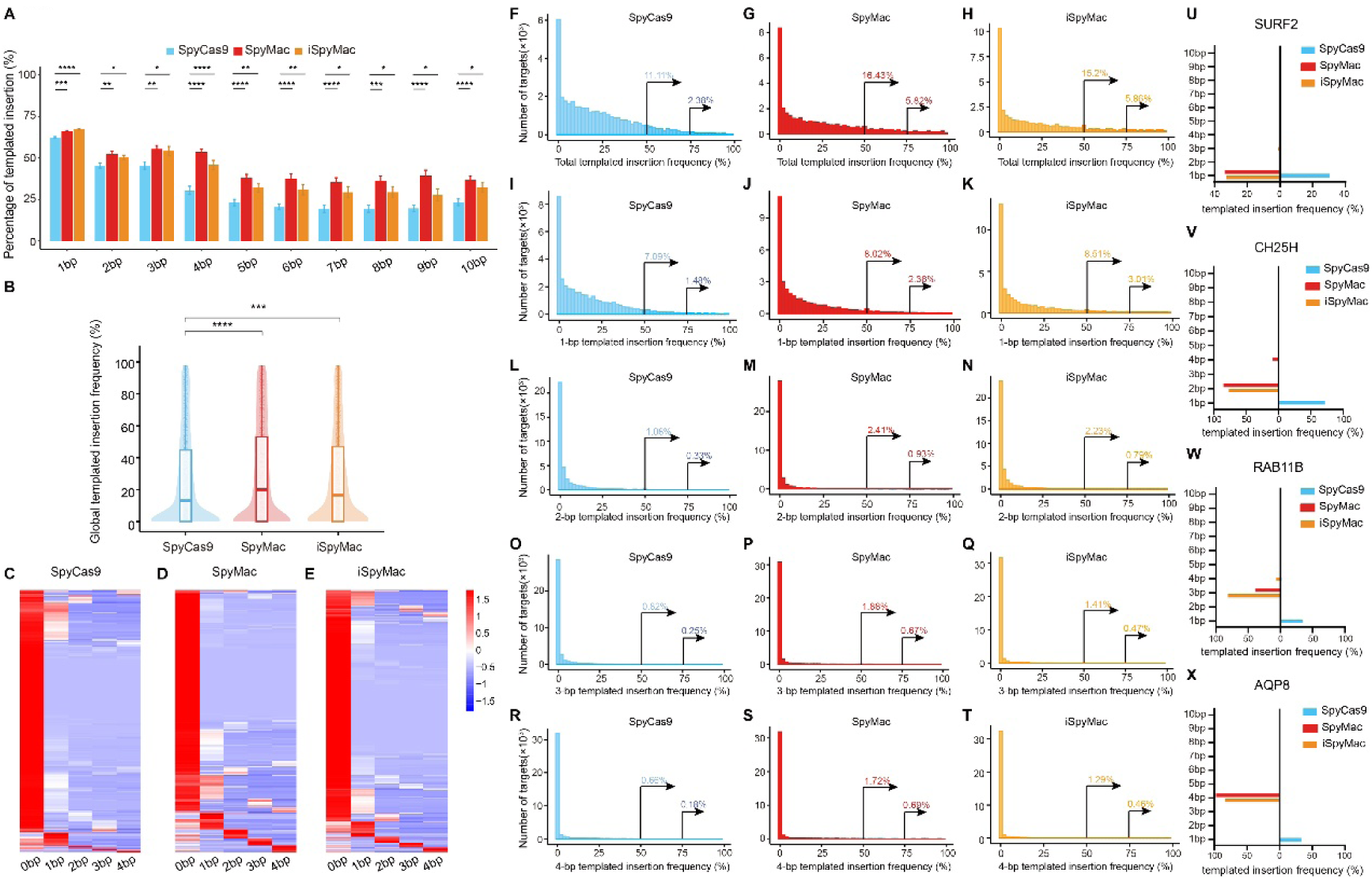
Precise templated insertions for A-rich PAM-recognizing Cas9 editors. (**A**) Percentages of templated insertions for SpyCas9, SpyMac, and iSpyMac across 37,266 targeting sites. Percentage of each of 1-10bp templated insertions is calculated as a ratio to the corresponding 1-10bp total insertions (templated insertions plus random insertions). (**B**) Global templated insertion frequency across targeting sites. Violin plots summarize per-target templated insertion frequencies; box plots indicate median and interquartile range. (**C-E**) Heatmaps of 1-4 bp templated insertions for SpyCas9 (C), SpyMac (D), and iSpyMac (E) across all targeting sites, highlighting distinct distributions of predicted overhang lengths among editors. Note the prevalence of 1-4bp templated insertion for A-rich PAM-recognizing editors. (**F-H**) Distributions of total templated insertion frequency for SpyCas9 (F), SpyMac (G), and iSpyMac (H). Percentages indicate the fraction of targeting sites exceeding the indicated templated insertion-frequency thresholds. (**I-T**) Distributions of 1-4bp templated insertion frequencies for SpyCas9 (I,L,O,R), SpyMac (J,M,P,S), and iSpyMac (K,N,Q,T). (**U-X**) Representative loci of SURF2 (U), CH25H (V), RAB11B (W), and AQP8 (X) illustrating distinct templated insertion profiles. For comparisons, statistical significance was assessed in R using an unpaired, two-sided Wilcoxon rank-sum test (Mann-Whitney U test); *P* values are denoted by asterisks as shown; \**p* ≤ 0.05, \*\**p* ≤ 0.01, \*\*\**p* ≤ 0.001, and \*\*\*\**p* ≤ 0.0001.

We then compared global templated insertion frequencies of 37,266 sites between A-rich and G-rich PAM-recognizing Cas9 editors and found that the A-rich PAM-recognizing Cas9 editors (SpyMac and iSpyMac) exhibited significantly higher templated insertion frequencies (Fig. 1B). Heatmap indicated a significantly higher proportion of targeting sites with 1-4bp dominant templated insertions for SpyMac and iSpyMac than for SpyCas9 (Fig. 1C-E). In addition, analyzing cleavage data[6,29] revealed pronounced staggered cleavage with long overhangs (Supplementary Fig. S5B-D). To further characterize insertions, we compared the distributions of templated insertion frequency for each of the 37,266 sites and found that 11.11%, 16.43%, and 15.2% of targeting sites have >50% templated insertions for SpyCas9, SpyMac, and iSpyMac, respectively (Fig. 1F-H). Analyzing the distributions of 1-4 bp templated insertion frequencies showed that the A-rich Cas9 editors have higher percentage of 1-4 bp than the G-rich SpyCas9 (Fig. 1I-T). Examples of 1-bp, 2-bp, 3-bp, and 4-bp templated insertional profiles are shown in Fig. 1U-X, respectively.

Finally, we analyzed random insertion patterns, which have long been recognized as a significant source of inaccuracy. We found that A-rich PAM-recognizing Cas9 editors exhibited significantly reduced frequencies of 1-10 bp random insertions in comparison to SpyCas9 (Fig. S5E). This is consistent with the enhanced precise insertions observed in A-rich PAM-recognizing Cas9 editors (Fig. 1A).

### Distinct types of deletions in A-rich and G-rich PAM-recognizing Cas9 editors

To comprehensively characterize deletion patterns of the 37,266 sites, we systematically analyzed the types and distributions of deletions induced by both A- and G-rich PAM-recognizing Cas9 editors. Deletions were classified based on molecular mechanisms into two primary categories: microhomology-mediated end joining (MMEJ) and non-MMEJ deletions. The non-MMEJ deletions were further categorized into three subtypes according to their positional characteristics relative to the cleavage site: bidirectional deletions covering both directions from the cleavage site, PAM-distal unidirectional deletions, and PAM-proximal unidirectional deletions (Fig. 2A). Our analyses revealed remarkable differences in deletion profiles between A-rich and G-rich editors, with MMEJ deletions accounting for 56.58%, 32.45%, 37.75% of the total deletions for SpyCas9, SpyMac, and iSpyMac, respectively (Fig. 2A). We next analyzed the microhomology usage[30] and found that the deletion frequencies decreased sharply as the size of microhomology increases (Fig. 2B-D). Linear regression analyses revealed that A-rich PAM-recognizing Cas9 editors (SpyMac and iSpyMac) have steeper slopes than SpyCas9 (Fig. 2E), indicating enhanced dependence on short sizes of microhomology.

**Figure 2.**
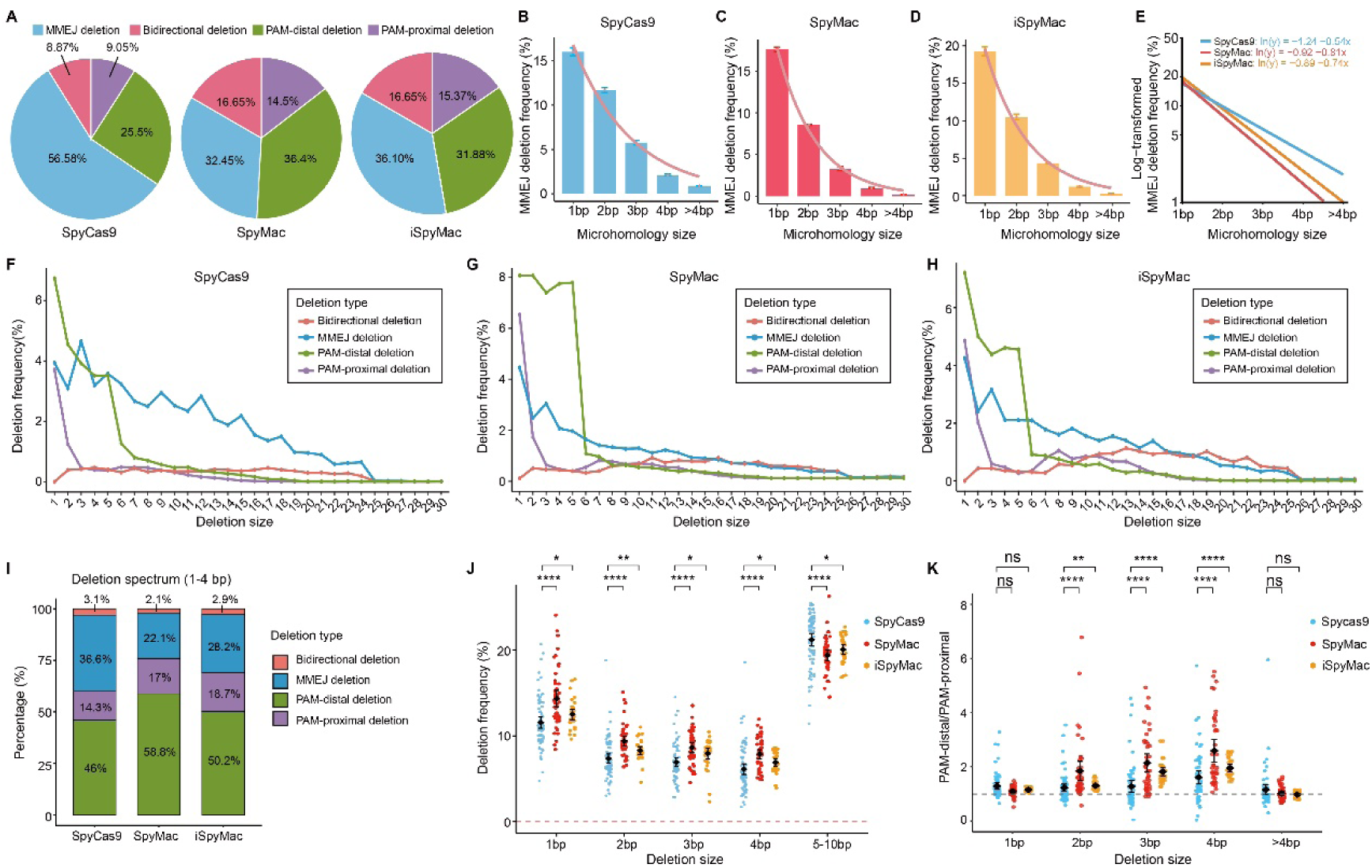
Deletion profiling for A-rich and G-rich PAM-recognizing Cas9 editors. (**A**) Distribution of microhomology-mediated end joining (MMEJ) deletions and non-MMEJ (bidirectional, PAM-distal, and PAM-proximal) deletions for SpyCas9, SpyMac, and iSpyMac. PAM-distal and PAM-proximal deletions were classified relative to the cleavage site, based on whether the deleted sequence lies on the PAM-distal or PAM-proximal side, respectively. (**B-D**) Microhomology size in MMEJ deletions for SpyCas9 (B), SpyMac (C), and iSpyMac (D). Fitted curves summarize the dependence on the microhomology size. (**E**) Log-transformed representation of the microhomology-size dependence with linear fittings indicated for each editor. (**F-H**) Deletion length spectra (1-30 bp) of the four deletion categories for SpyCas9 (F), SpyMac (G), and iSpyMac (H). (**I**) Deletion profiles for SpyCas9, SpyMac, and iSpyMac. (**J**) Comparison of normalized frequencies of small deletions of SpyCas9, SpyMac, and iSpyMac. (**K**) Ratio of PAM-distal to PAM-proximal deletions for SpyCas9, SpyMac, and iSpyMac. *P* values were computed in R using an unpaired, two-sided Wilcoxon rank-sum test (Mann-Whitney U test); \**p* ≤ 0.05, \*\**p* ≤ 0.01, and \*\*\*\**p* ≤ 0.0001; ns, not significant.

We then examined the deletion profiles across different lengths (1-30 bp) and found that the A-rich PAM-recognizing Cas9 editors exhibited a more pronounced enrichment of small deletions at PAM distal side than the G-rich SpyCas9 (Fig. 2F-H). Quantification analyses revealed that 1-4 bp PAM-distal deletions, which are resulted from nuclease digestion of single-stranded overhangs of staggered cleavage[11], occurred at significantly higher frequencies in both SpyMac and iSpyMac than in SpyCas9, consistent with more staggered cleavage for A-rich PAM editors (Fig. 2I). In addition, for each of the four small deletions (1-4 bp), SpyMac and iSpyMac have a significant higher deletion frequency than SpyCas9 (Fig. 2J). Finally, using the ratio of PAM-distal to PAM-proximal deletions as an asymmetry index, we found that A-rich PAM-recognizing Cas9 editors of SpyMac and iSpyMac have significant more 2-4bp bias toward PAM-distal deletions, indicating that the A-rich Cas9 editors generate more staggered 5’ ends than the G-rich SpyCas9 (Fig. 2K). All in all, these data demonstrated that the A-rich Cas9 editors are more precise than G-rich SpyCas9 for PAM-distal deletions.

### MMEJ deletions mediated by single nucleotide microhomology explain the predominant −4bp insertions

Comprehensive analyses of local sequence contexts revealed a DNA motif for templated insertions (Supplementary Fig. S6A-C). To solve the underlying mechanism, we first analyzed the effect of each of the four nucleotides at −4 position on templated insertions and found that, consistent with the DNA motif, nucleotides “A/T” at −4 position favor templated insertions of staggered cleavage, whereas nucleotides “G/C” at −4 position disfavor templated insertions for all three Cas9 editors (Fig. 3A). Intriguingly, we observed a significant decrease of templated insertions of nucleotides “A/T” at −4 position for A-rich PAM compared to G-rich PAM (Fig. 3A). In addition, with nucleotides “G/C” at −4 position, we observed a significant increase of templated insertions for the A-rich PAM compared to the G-rich PAM (Fig. 3A).

**Figure 3.**
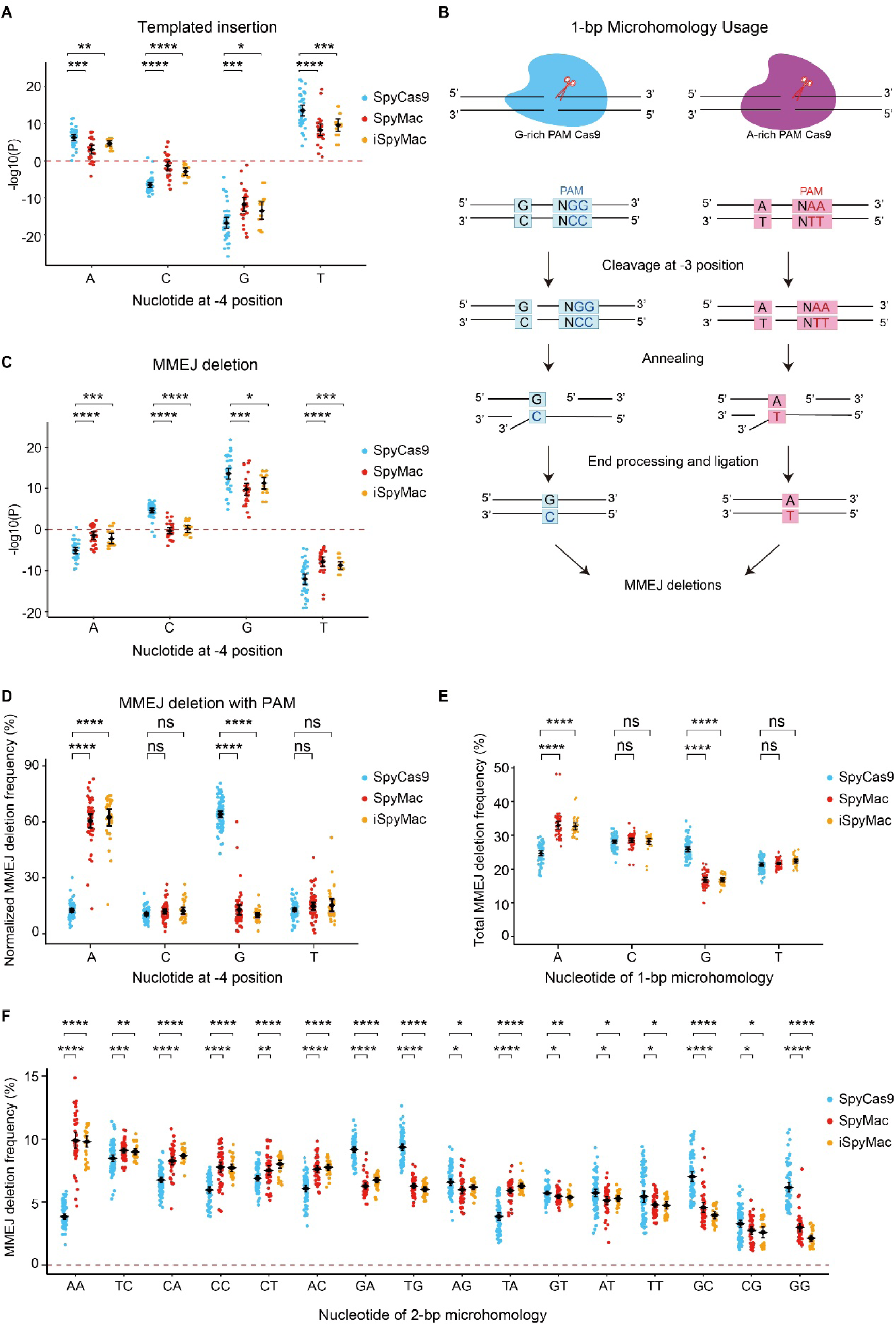
Mechanism for DNA motif of templated insertions. (**A**) Effect of the −4 nucleotides (numbered relative to PAM) on templated insertion of SpyCas9, SpyMac, and iSpyMac. (**B**) Model of MMEJ with single nucleotides at −4 annealing to the PAM nucleotides during end joining. Schematics contrast microhomology usage in G-rich PAM (NGG) versus A-rich PAM (NAA) contexts. (**C**) Effect of the −4 nucleotides on deletion. (**D**) Normalized deletion frequency between −4 nucleotides and PAM sequences. (**E**) Frequencies of MMEJ deletions mediated by 1-bp microhomologies (A, C, G or T). (**F**) Frequencies of MMEJ deletions mediated by 2-bp microhomologies across all 16 dinucleotide combinations. *P* values were computed in R using an unpaired, two-sided Wilcoxon rank-sum test (Mann-Whitney U test); \**p* ≤ 0.05, \*\**p* ≤ 0.01, \*\*\**p* ≤ 0.001, and \*\*\*\**p* ≤ 0.0001; ns, not significant.

We reasoned that the −4 “G” may be a 1-bp microhomology with the G-rich PAM, leading to MMEJ deletions, and thus disfavoring 1bp insertions (Fig. 3B). To this end, we analyzed paired editing outcomes of A-rich and G-rich PAMs for each of the 37,266 targeting sites. Remarkably, we observed a significant increase of deletions for −4 “A” with the A-rich PAM of SpyMac and iSpyCas9 in comparison to the G-rich PAM of SpyCas9 (Fig. 3C and Supplementary Fig. S6D-F). In addition, we also observed a significant increase of deletions for −4 “G” with the G-rich PAM of SpyCas9 compared to the A-rich PAM (Fig. 3C). Moreover, we specifically analyzed MMEJ deletions between −4 nucleotide and PAM and observed striking increases between “A” and NAA or −4 “G” and NGG, with no significant alternation for internal control of −4 “C” or “T” (Fig. 3D). These observations explained the long-standing mystery of −4 “G” in the DNA motif disfavors 1bp insertions of SpyCas9 editing outcomes[5,6,8,25–27,30].

We next examined total MMEJ deletion events mediated by 1-bp microhomology and found a significant increase of MMEJ deletions mediated by the single nucleotide “A” for the A-rich Cas9 editors compared with SpyCas9 (Fig. 3E). Conversely, there is a significant decrease of MMEJ deletions mediated by the single nucleotide “G” for the A-rich Cas9 editors compared with SpyCas9 (Fig. 3E). As internal controls, there is no significant alteration of MMEJ deletion frequency for the −4 position of “C” or “T” between these G-rich and A-rich PAM editors. Together, these observations clearly showed that MMEJ deletions mediated by single-nucleotide homologies play a key role in shaping sequence-context patterns for predicting Cas9 editing outcomes.

Finally, we comprehensively analyzed 2-bp microhomology usage of all of the 16 possible dinucleotide combinations in MMEJ deletions (Fig. 3F). We observed a remarkable decrease of MMEJ deletions from dinucleotides “AA” to “GG” for A-rich PAM editors (Fig. 3F). In addition, “AA” dinucleotides showed remarkably enhanced MMEJ deletions for the A-rich PAM of SpyMac and iSpyMac. By contrast, “GG” dinucleotides showed remarkably enhanced MMEJ deletions for the G-rich PAM of SpyCas9 (Fig. 3F). Altogether, these data suggest that the PAM sequences play a crucial role in Cas9 precision editing.

### PolQ plays a role in random insertions

To investigate the function of PolQ in precision editing, we performed large-scale profiling under PolQ-deficient conditions. We first generated six PolQ-deficient cell lines, with two for each deletion of its entire coding region (ΔPolQ), of polymerase domain (ΔPol), and of helicase domain (ΔHel) through CRISPR screening of 356, 306, and 319 Cas9-expressing single-cell HEK293T clones, respectively (Fig. 4A). We then transduced the sgRNA libraries into each of the six PolQ-deficient single-cell clones and analyzed ∼180 million sets of editing outcomes and found a high inter-clonal consistency in editing efficiency (Supplementary Fig. S7A-D). We analyzed the Shannon index and its dependency on the nucleotide compositions and found they are largely unaffected (Supplementary Fig. S7E-S).

**Figure 4.**
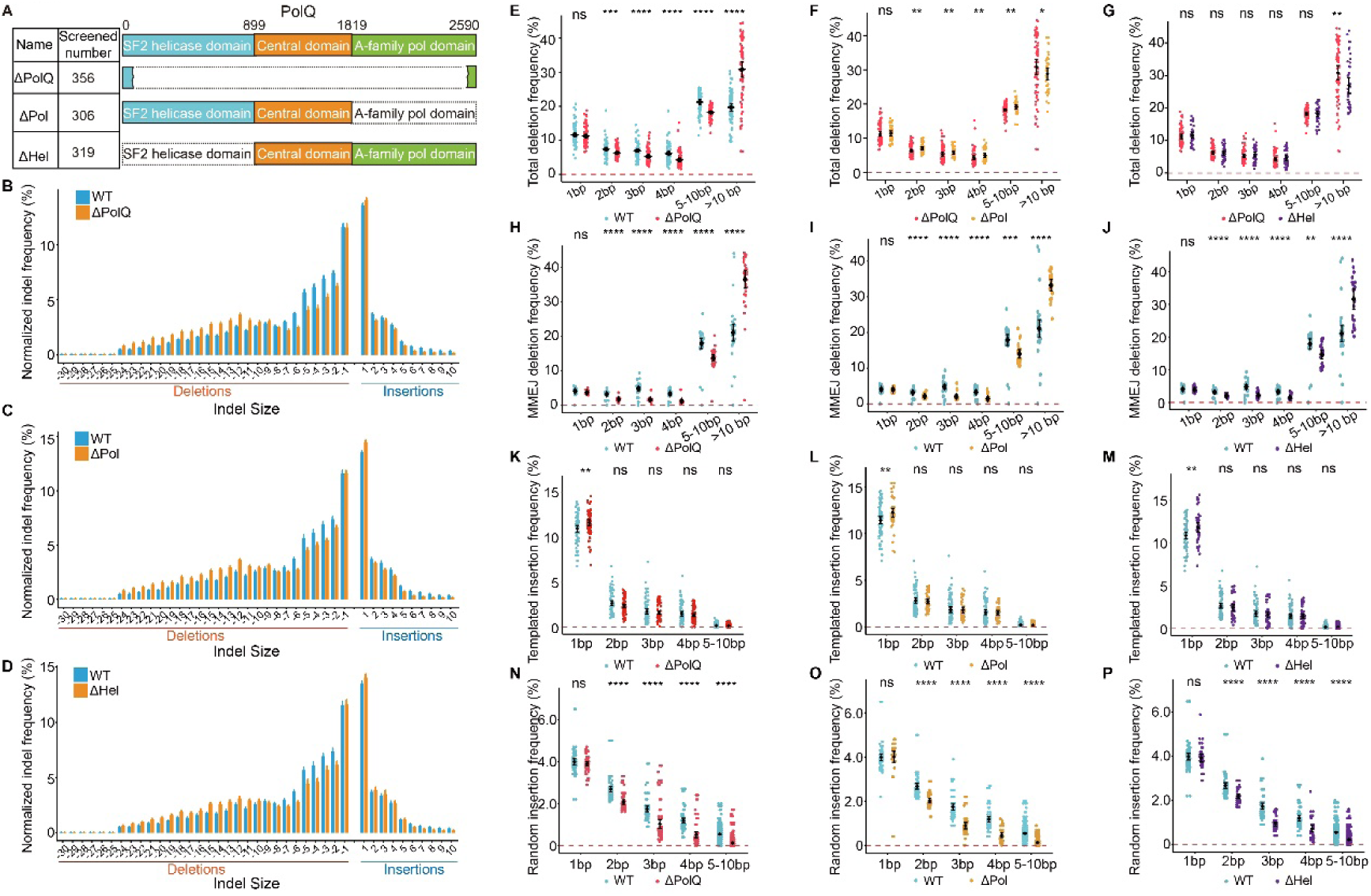
PolQ deficiency decreases random insertions. (**A**) Schematic of POLQ domain organization and CRISPR strategies used to generate single-cell HEK293T clones with deletion of the entire POLQ coding region (ΔPolQ), the polymerase domain (ΔPol), or the helicase domain (ΔHel). (**B-D**) Distribution of normalized indel frequency of 37,266 targeting sites in ΔPolQ (B), ΔPol (C), or ΔHel (D) compared with WT cells. (**E-G**) Effects of PolQ deficiencies on total deletion frequencies. (**H-J**) Effects of PolQ deficiencies on MMEJ deletion frequencies. (**K-M**) Effects of PolQ deficiencies on templated insertion frequencies. (**N-P**) Effects of PolQ deficiencies on random insertion frequencies. *P* values were calculated in R using an unpaired, two-sided Wilcoxon rank-sum test; \**p* ≤ 0.05, \*\**p* ≤ 0.01, \*\*\**p* ≤ 0.001, and \*\*\*\**p* ≤ 0.0001; ns, not significant.

We next analyzed indel profiles and found widespread shifts in indel landscapes upon PolQ deficiencies (Fig. 4B-D). Specifically, there is a significant decrease of small deletions (2-10 bp) and a significant increase of larger deletions (>10 bp) upon PolQ deletion (Fig. 4E). By contrast, there is no significant change of 1-bp deletions (Fig. 4E), consistent with that 1-bp deletions occur by the PolQ-independent non-homologous end joining (NHEJ) pathway[11]. To dissect the domain function of PolQ, we first deleted the PolQ polymerase domain (Fig. 4A) and found that there is a significant increase of 2-10bp deletions and a significant decrease of >10bp deletions in comparison with the entire PolQ deletion (Fig. 4F). We then deleted the helicase domain and found that only >10 bp deletions are significantly decreased, suggesting that the helicase domain may function in resolving the large-sized microhomology (Fig. 4G). Furthermore, the deletion profiles and microhomology usage appear to be similar (Supplementary Fig. S8A-F). Finally, we analyzed MMEJ deletions and found that there are a significant decrease of small deletions (2-10 bp) and a significant increase of larger deletions (>10 bp) upon PolQ deficiencies (Fig. 4H-J and Supplementary Fig. S8G-L).

We then analyzed templated insertions and found that there is a significant increase of 1bp, but not 2-10bp, templated insertions upon PolQ deficiencies (Fig. 4K-M). Finally, we analyzed random insertions and found that there is no significant change of 1bp random insertions upon PolQ deficiencies, consistent with that 1bp random insertions are generated by NHEJ (Fig. 4N-P and Supplementary Fig. S8M-R). Interestingly, we found a remarkable and consistent decrease of frequencies of 2-10bp random insertions upon PolQ deficiencies, suggesting that PolQ plays a crucial role in 2-10bp random insertions (Fig. 4N-P and Supplementary Fig. S8M-R).

### CRIfuser predicts genome-wide editing outcomes with staggered cleavages

To accurately estimate genome-wide staggered cleavage and subsequent repair outcomes, we developed CRISPR diffuser (CRIfuser), a discrete-space diffusion-based machine learning framework designed to predict editing profiles from ligations of the two processed DSB ends at any given cleavage site (Fig. 5A and Supplementary Note S1). To this end, we used the horizontal axis (*x*) to represent the linear sequence of the PAM-distal ligation end and the vertical axis (*y*) to represent that of the PAM-proximal ligation end. Thus, any base-pair sequence upstream of the −3 position of PAM in the *y* axis represents the precise and predictable templated insertions resulting from the fill-in of the 5’ overhang of PAM-proximal DSB end of staggered cleavage; by contrast, all previous predicting models do not even consider precise and predictable templated insertions from staggered or cohesive Cas9 cleavages. In addition, we designed a new loss function tailored for training the two-dimensional diffusion model for predicting CRISPR editing profiles, which significantly improved its convergence and accuracy. Finally, our diffusion model accounts for MMEJ precision deletions, which are bidirectional by definition, and MMEJ-independent deletions, which include unidirectional deletions (both PAM-distal and PAM-proximal) and random bidirectional deletions (Fig. 5A).

**Figure 5.**
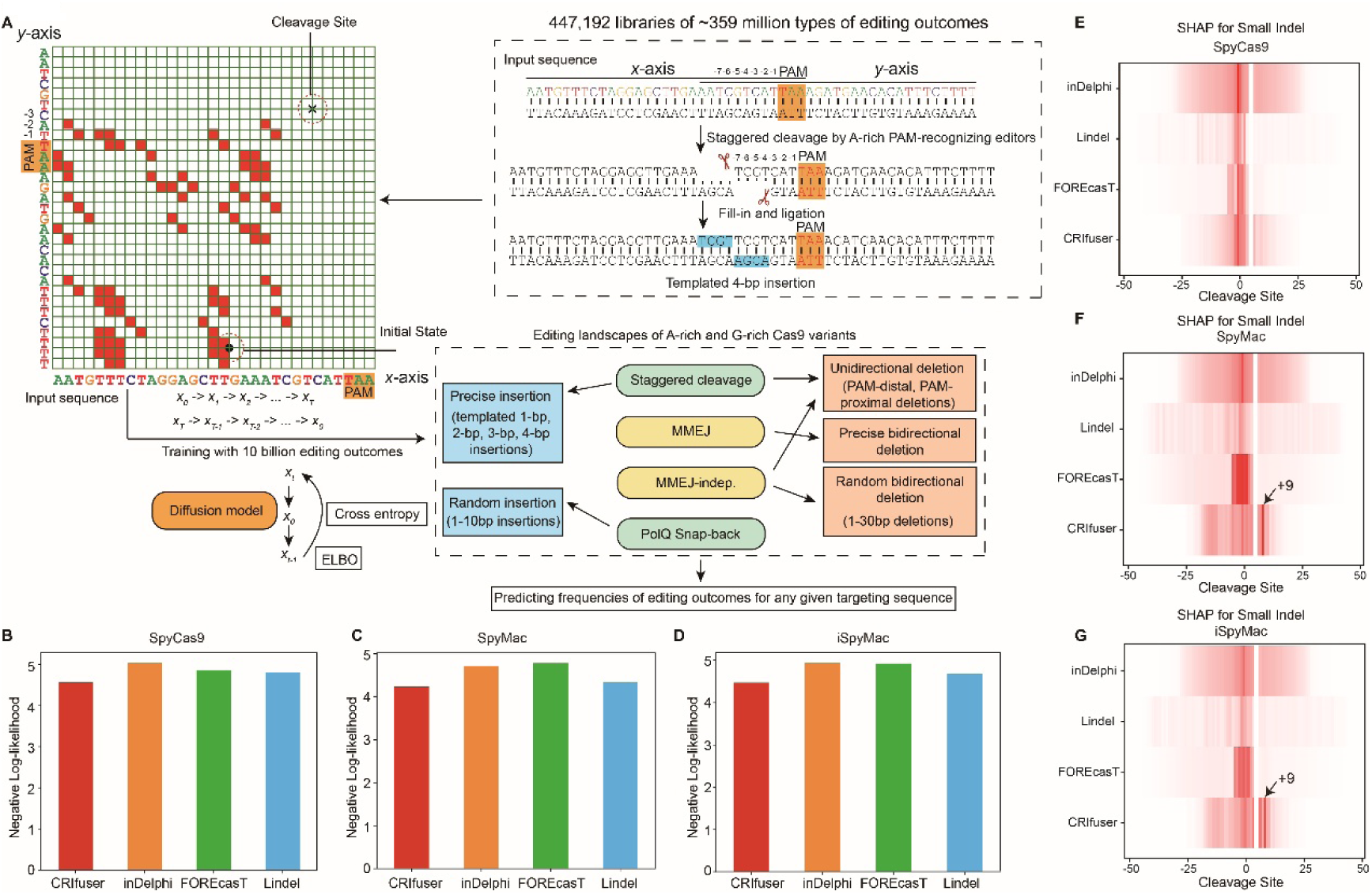
CRIfuser predicts genome-wide editing outcomes generated by staggered cleavages. (**A**) Overview of the discrete diffusion model of CRIfuser. Editing outcomes are represented as a two-dimensional distribution of ligation junction positions. Sequence context of cleavage site is used as input, and the discrete diffusion model is trained to predict repair profiles, including templated insertions, MMEJ deletions, and non-MMEJ deletions (bidirectional deletions, PAM-distal and PAM-proximal deletions). (**B-D**) Benchmark of CRIfuser with inDelphi, FOREcasT, and Lindel for SpyCas9 (B), SpyMac (C), and iSpyMac (D). (**E-G**) Heatmaps showing SHAP (SHapley Additive exPlanations) importance of bases around cleavage site in predicting editing outcomes of small indels for SpyCas9 (E), SpyMac (F), and iSpyMac (G). Note the strong signals at the +9 position predicted by CRIfuser for SpyMac and iSpyMac but not SpyCas9.

To establish a benchmark, we compared CRIfuser with inDelphi, FOREcasT, and Lindel (Supplementary Note S2). We evaluated model performance using Negative Log-Likelihood (NLL) loss between the predicted profiles and the observed editing outcomes for SpyCas9 (Fig. 5B), SpyMac (Fig. 5C), and iSpyMac (Fig. 5D) and found that CRIfuser achieved the best performance (Fig. 5B-D and Supplementary Fig. S9A-C). We then employed SHAP (SHapley Additive exPlanations)[31] to interpret model predictions and to evaluate the importance of sequence contexts around the cleavage site. Interestingly, we found that the +9 position relative to the cleavage site is a critical determinant for the editing outcomes of SpyMac and iSpyMac but not of SpyCas9 (Fig. 5E-G and Supplementary Fig. S9D-L). Finally, to facilitate broad accessibility, we developed an interactive web application for editing outcome prediction based on the CRIfuser standalone software.

## Discussion

The protospacer adjacent motif (PAM) is a fundamental determinant of targeting specificity in the CRISPR/Cas systems. PAM recognition licenses Cas9 target binding, DNA strand separation, and R-loop formation, thereby constraining editable loci and reducing off-target cleavage at PAM-incompatible sites[2,13–15]. In this study, we conducted a systematic comparison of two A-rich PAM-recognizing Cas9 editors (SpyMac and iSpyMac) with the canonical G-rich SpyCas9. We mapped ∼half-million sets of ∼10 billion editing outcomes from ∼37 thousand protospacers and found that the A-rich SpyMac and iSpyMac exhibited superior editing precision. In particular, these A-rich editors showed significantly enhanced efficiency for templated insertions from their staggered cleavage and thus resulting in predictable and precision editing. Thus, PAM recognition is not merely a targeting constraint but an active parameter in the design of high-precision editing strategies, influencing staggered cleavage probability, repair pathway engagement, and overall editing accuracy. Finally, we analyzed editing profiles under PolQ-deficient conditions and found its crucial role in precise deletions and random insertions.

Although SpyCas9 has been widely characterized as generating blunt DSB ends 3bp upstream of PAM[2], accumulating biochemical and genetic analyses demonstrate that Cas9 cleavage is frequently staggered, producing 1-3 nucleotide short 5′ overhangs due to asymmetric activity of the HNH and RuvC nuclease domains[5,6,26]. Specifically, the HNH domain cleaves the target strand at the exact −3 position upstream of PAM, whereas the cleavage by the RuvC domain on the non-target strand is relaxed and flexible[8,32,33]. While the formation of 1-bp 5’ overhangs by SpyCas9 is well-established[4,5,25], a recent study proposed that multi-nucleotide insertions can be explained by Cas9 target recleavage[34]. The study tested only one target sequence that happens to have a dinucleotide “TT” at the −4 and −5 positions upstream of PAM[34], while *in-vitro* cleavage on multiple targeting sequences with no repair, and thus no recleavage, clearly showed direct staggered DSB ends with flexible overhang profiles (Supplementary Fig. S5B-D)[6,8]. By leveraging custom algorithms designed to accurately analyze templated insertions, we demonstrated that the frequency of longer templated insertions, in particular the preponderance of 4bp from staggered cleavage, significantly exceeds what would be expected from stochastic chances (Fig. 1 and Supplementary Fig. S5A-D). This high propensity, observed across both G-rich and A-rich PAM-recognizing Cas9 editors, strongly suggests that these longer templated insertions represent genuine editing outcomes derived from inherent staggered cleavage profiles. In particular, the RuvC domain exhibits positional flexibility on the non-target strand, leading to heterogeneous cleavage sites and non-blunt DNA ends[5,6,9]. These staggered cleavage products have important functional consequences for repair pathway choice, as non-homologous end joining (NHEJ) preferentially fills in 5′ overhangs prior to ligation, resulting in templated insertions with tandem head-to-tail duplications, whereas alternative pathways such as microhomology-mediated end joining (MMEJ) contribute to more diverse deletion outcomes[11,12].

The discovery of Cas9 staggered cleavage mechanism has had profound impacts on the development of precision editing technologies and clinical applications[2,5]. For example, utilizing destabilization induced by relaxed RuvC nicks further upstream of PAM on the noncomplementary strand has led to the development of next-generation prime editor with strikingly low indel errors[32,33]. In addition, overhang-dependent base replication of Cas9 staggered cleavage enables successful corrections of a human muscular dystrophy founder mutation[35]. From an engineering perspective, PAM identity further modulates Cas9 binding stability, cleavage efficiency, and editing window properties, thereby influencing both editing efficiency and product distribution. Collectively, the interplay between PAM recognition, staggered cleavage asymmetry, and DNA repair pathway choice defines a tunable framework for optimizing editing precision and expanding the targeting scope of CRISPR-based technologies.

Repair outcomes exhibited a strong dependency on the protospacer sequence at the target site. For SpyCas9, an idiosyncratic DNA motif for templated insertions predicts that an “A/T” at the −4 position upstream of the PAM sequence promotes templated insertions, whereas a “G/C” at this position exerts an inhibitory effect[25,27,30,36]. However, the underlying mechanism has long remained mysterious. By employing hybrid Cas9 variants (SpyMac and iSpyMac) that recognize the NAA PAM with SpyCas9 for the paired NGG PAM as controls, we observed significant differences in DNA motifs for both templated insertions and MMEJ deletions. Remarkably, when an “A” was present at the −4 position, these A-rich PAM-recognizing editors were significantly less prone to induce templated insertions compared to SpyCas9; whereas a “G” at the same position, these variants exhibited a reduced propensity for deletions relative to their SpyCas9 counterpart (Fig. 3C). In particular, we investigated the deletions that occurred between the −4 nucleotide and the PAM sequence and found a remarkable nucleotide preference depending on the PAM specificity (Fig. 3D). Specifically, for the A-rich PAM-recognizing editors (SpyMac and iSpyMac), nucleotide “A” at −4 position yields significantly higher deletion frequencies than the other three nucleotides. Similarly, nucleotide “G” at −4 position mediates higher deletion efficiencies for SpyCas9, which recognizes a G-rich PAM. These intriguing observations of MMEJ deletions between −4 and PAM nucleotides clearly explained the DNA motif of templated insertions in addition to staggered cleavage[25,27,30,36]. Together, these observations provide strong evidence that microhomologies of not only typical 2-8bp previously known[37] but also even 1bp mediate MMEJ deletions.

Finally, the Polθ polymerase (encoded by PolQ) establishes a prominent role in alternative end joining such as MMEJ or theta-mediated end joining (TMEJ)[38–40], yet it is remarkable that substantial reduction of random insertions was observed under PolQ-deficient conditions (Fig. 4N-P and Supplementary Fig. S8M-R). This phenomenon could be rationalized by the synthesis-dependent microhomology-mediated end-joining (SD-MMEJ), which is associated with Polθ activity[41]. Mechanistically, Polθ facilitates alternative end-joining via its capacity to mediate both *in cis* (via loop-out and snap-back mechanisms) and *in trans* (via base-pairing from opposing resected single strand) insertions of stochastic flanking DNA sequences into cleavage sites[42–44]. Consequently, the abrogation of PolQ leads to the observed decrease in random insertions[38]. Conversely, the concurrent increase in 1-bp templated insertions can be attributed to pathway competition between cNHEJ and MMEJ. As PolQ deficiency impairs MMEJ, DSB repair is presumably channeled toward the canonical non-homologous end-joining (cNHEJ) pathway, which inherently favors the generation of 1-bp templated insertions upon Cas9 staggered cleavage[11].

In conclusion, our data suggest PAM identity as an important determinant of editing outcomes and highlight the A-rich PAM-recognizing editors not only expanding targeting sequence space, but also as promising tools for precision genome editing. By preferentially generating more predictable repair products through enhanced staggered cleavage and templated insertion, these A-rich PAM-recognizing editors offer improved control for gene editing.

## Acknowledgements

We are grateful for discussions from all members of our laboratory.

## Author contributions

Q.W. conceived the research. X.S., C.L., J.L., and Y.L. performed experiments. X.S., C.L., and J.L. analyzed data. J.L. developed the AI model. X.S., C.L., J.L., and Q.W. wrote the manuscript. All authors have read and approved the final manuscript.

## Conflict of interest

None declared.

## Funding

This study was supported by grants to Q.W. from the National Natural Science Foundation of China (32330016 and 32670802) and the National Key R&D Program of China (2022YFC3400200).

## Data availability

The data produced in this study are openly available in GSA under accession number HRA018811. Both the web tool and the open-source software are available at https://hf.co/spaces/ljw20180420/CRISPR_AI and GitHub https://github.com/ljw20180420/CRISPR_AI.

Any additional information required to reanalyze the data reported in this paper is available from the lead contact upon request.

## Notes

### Competing Interest Statement

The authors have declared no competing interest.

