## Supplementary figures and notes for "Large scale CRISPR genome engineering reveal principles of single nucleotide microhomology in precision editing"

Figure S1

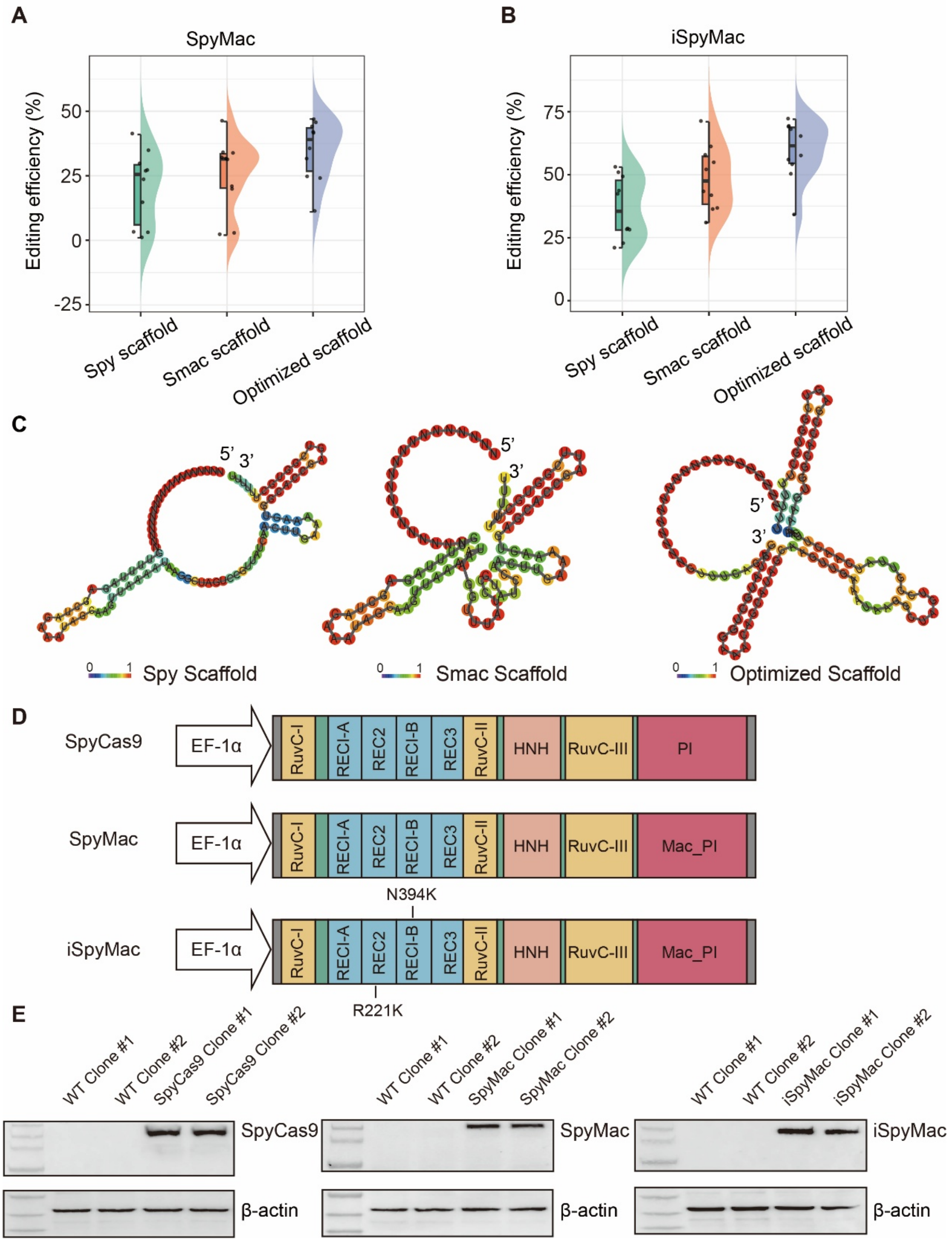


**Figure S1.** sgRNA scaffold optimization for A-rich PAM-recognizing Cas9 editors. (**A,B**) Editing efficiencies of ten targeting sites obtained with Spy, Smac, and optimized sgRNA scaffolds for SpyMac (A) and iSpyMac (B). (**C**) Predicted secondary structures of the three sgRNA scaffolds. (**D**) Schematics of SpyCas9, SpyMac, and iSpyMac constructs. (**E**) Western blot analyses showing expression levels for SpyCas9, SpyMac, and iSpyMac in stable single-cell clones.

Figure S2

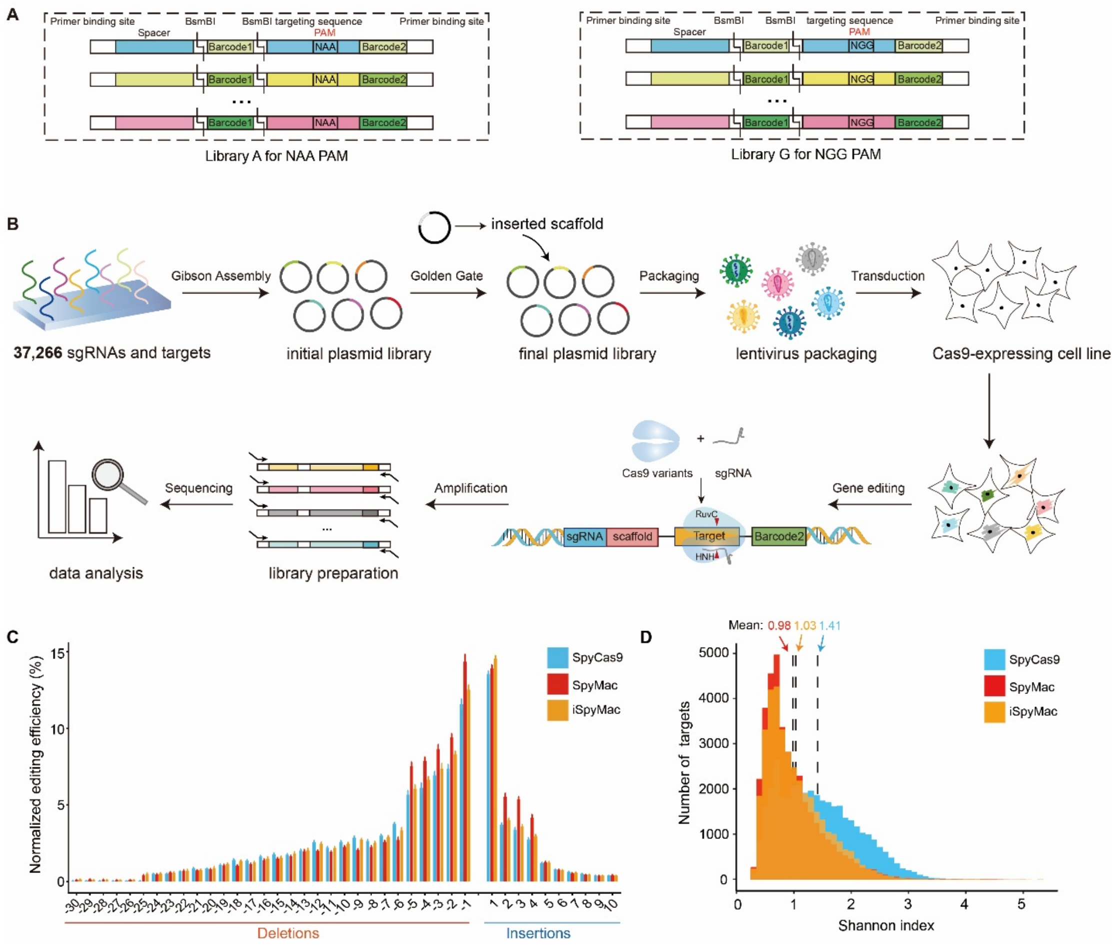


**Figure S2.** High-throughput profiling for A-rich PAM-recognizing Cas9 editors. (**A**) Design of paired high-throughput oligonucleotide libraries comprising 37,266 targeting sites. Library A contains targets bearing an NAA PAM, whereas Library G contains the identical set of protospacers paired with the NGG PAM controls. Each oligonucleotide contains a spacer and its cognate target sequence separated by a 20-bp internal barcode (barcode1) flanked by *Bsm*BI sites, and includes a second downstream barcode (barcode2) for amplicon identification. (**B**) Workflow for library construction and editing profiling. Array-synthesized oligonucleotide pools were amplified and cloned into a lentiviral backbone to generate an initial plasmid library, followed by insertion of sgRNA scaffold by Golden Gate cloning to generate the final library. Lentiviral libraries were transduced into stable SpyCas9-, SpyMac-, or iSpyMac- expressing HEK293T single-cell clones, followed by amplicon preparation and high-throughput sequencing to quantify editing outcomes. (**C**) Length distribution of insertions and deletions across 37,266 targeting sites edited by SpyCas9, SpyMac, and iSpyMac, shown as normalized frequencies for each indel size class. Note that 1-4 bp insertions and 1-5 bp deletions are abundant editing outcomes of the three Cas9 editors. (**D**) Distribution of per-target editing outcome diversity quantified by Shannon index for SpyCas9, SpyMac, and iSpyMac; dashed vertical lines indicate the mean Shannon index for each editor.

Note that both the A-rich SpyMac (Mean = 0.98) and iSpyMac (Mean = 1.03) have a lower Shannon index than the G-rich SpyCas9 (Mean = 1.41).

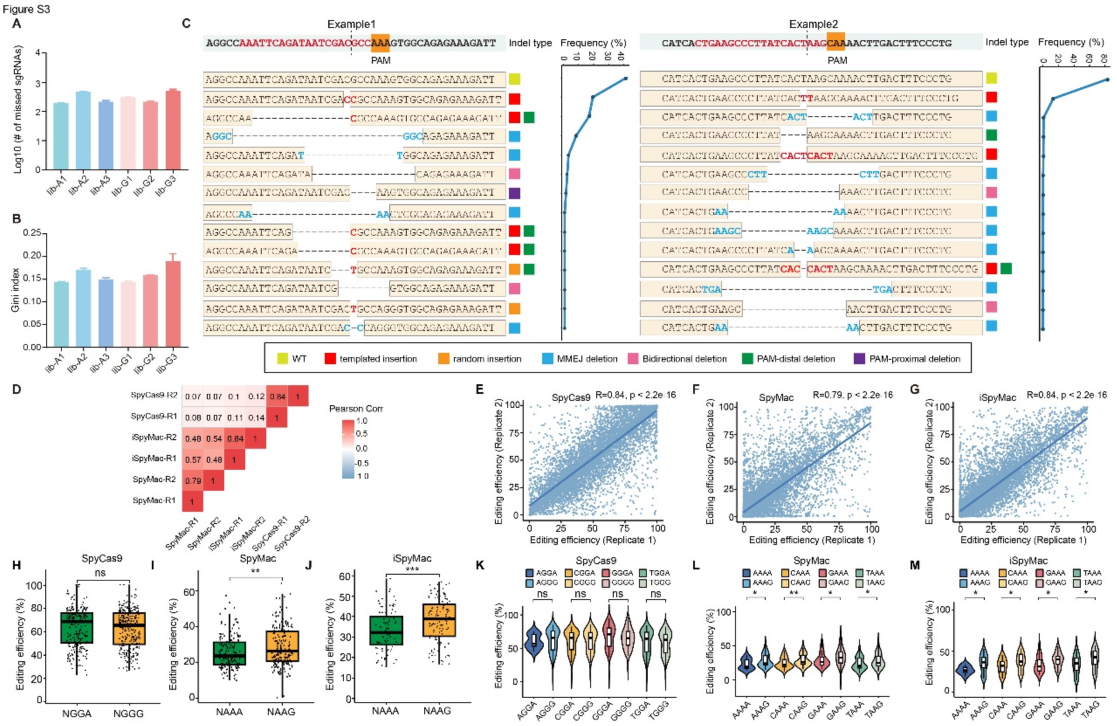


**Figure S3.** Quality control of high-throughput sgRNA libraries and analysis of editing outcomes. (**A**) Library coverage assessment for each constructed library, each of the six libraries has a high coverage of sgRNAs, shown as the number of missing sgRNAs (log10 scale). (**B**) Uniformity analyses of sgRNA representation across libraries quantified by the Gini index. The Gini index for our libraries was approximately 0.15, indicating a highly uniform distribution of sgRNAs. (**C**) Representative examples of editing outcomes, including insertions, deletions, and indels. Note that templated insertions of staggered cleavage result in tandem or head-to-tail duplications (“CC”, “TT”, and “CACTCACT”) of nucleotides immediately upstream of -3 position of PAM. Nucleotides of junctional microhomologies are highlighted in blue. (**D**) Heatmap of Pearson correlation showing the robust inter-replicate reproducibility of editing efficiencies for each Cas9 editor (SpyCas9, SpyMac, and iSpyMac). (**E-G**) Scatter plots showing the concordance of editing efficiencies for SpyCas9 (E), SpyMac (F), and iSpyMac (G) between biological replicates, with Pearson correlation coefficients and *P* values indicated. (**H-M**) Editing efficiencies across all eight NAAR PAM subtypes via a global analysis of the editing outcomes of 37,266 targeting sites: SpyCas9 NGGA versus NGGG (H,K), SpyMac NAAA versus NAAG (I,L), and iSpyMac NAAA versus NAAG (J,M), highlighting preferential activity on NAAG relative to NAAA and indicating that the fourth nucleotide of PAM is important for the two A-rich Cas9 editors of SpyMac and iSpyMac. Statistical significance was assessed in R using an unpaired, two-sided Wilcoxon rank-sum test (Mann-Whitney U test); *P* values are denoted by asterisks as shown; $*p \leq 0.05$ and $**p \leq 0.01$; ns, not significant.

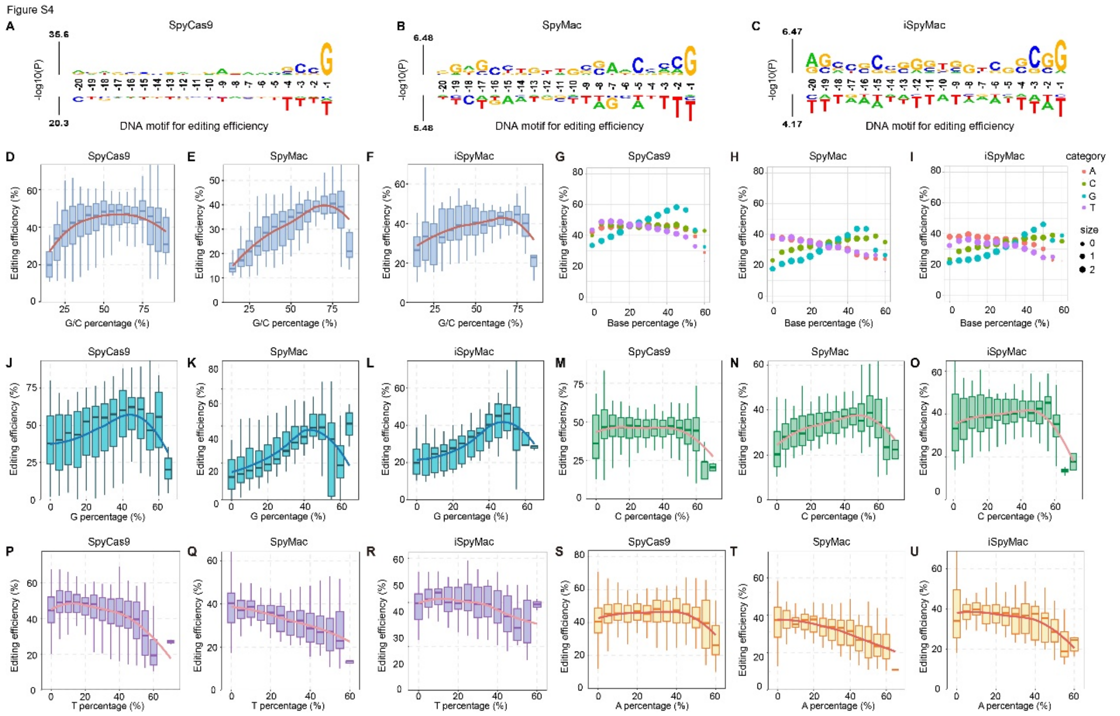


**Figure S4.** A-rich and G-rich PAM-recognizing Cas9 editors exhibit distinct sequence-context dependencies for editing efficiency. (**A-C**) kpLogo analysis of protospacer-sequence features associated with editing efficiency for SpyCas9 (A), SpyMac (B), and iSpyMac (C). A-rich PAM-recognizing Cas9 editors of SpyMac and iSpyMac exhibit positional dependencies on both the seed and non-seed regions; whereas only the seed region upstream of PAM is important for SpyCas9 with guanine (G) at the -1 position showing high editing efficiency. Enriched nucleotides at each protospacer position are shown above the axis and depleted nucleotides below the axis. Positions are numbered relative to the PAM, with -20 to -1 corresponding to the 20-nt protospacer sequence immediately upstream of PAM, and with heights indicating statistical significance (-log10 P). (**D-F**) Dependence of editing efficiency on protospacer G/C content for SpyCas9 (D), SpyMac (E), and iSpyMac (F). Targeting sites were grouped by protospacer G/C percentage and summarized as box plots, with a fitted trend line overlaid. (**G-I**) Nucleotide-composition dependency of editing efficiency for SpyCas9 (G), SpyMac (H), and iSpyMac (I). (**J-L**) Editing efficiency as a function of protospacer G percentage for SpyCas9 (J), SpyMac (K), and iSpyMac (L). (**M-O**) Editing efficiency as a function of protospacer C percentage for SpyCas9 (M), SpyMac (N), and iSpyMac (O). (**P-R**) Editing efficiency as a function of protospacer T percentage for SpyCas9 (P), SpyMac (Q) and iSpyMac (R). (**S-U**) Editing efficiency as a function of protospacer A percentage for SpyCas9 (S), SpyMac (T), and iSpyMac (U).

Figure S5

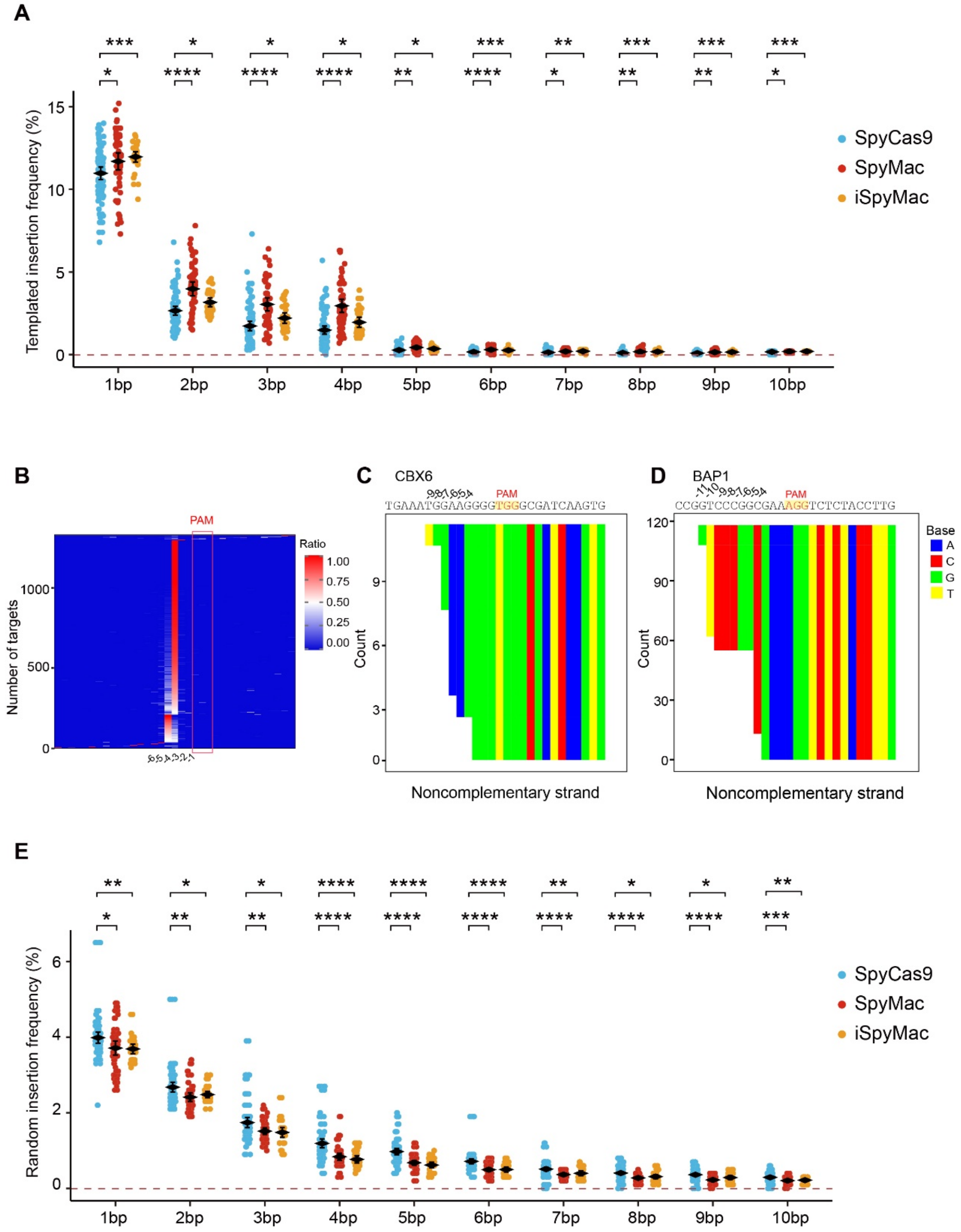


**Figure S5.** A-rich PAM-recognizing Cas9 editors exhibit higher templated insertions and lower random insertions. (**A**) Frequencies of 1-10bp templated insertions in total edited reads for SpyCas9, SpyMac, and iSpyMac. Statistical significance between editors for each length is indicated above the plot. (**B-D**) Heatmaps (B) and examples (C and D) of *in-vitro* cleavage profiles of 1,418 targeting sites[6,29], showing pronounced staggered cleavage on the noncomplementary strand. (**E**) Frequencies of 1-10bp random (non-templated) insertions in total edited reads for SpyCas9, SpyMac, and iSpyMac. *P* values were calculated in R using an unpaired, two-sided Wilcoxon rank-sum test; $^{*}p \leq 0.05$, $^{**}p \leq 0.01$, $^{***}p \leq 0.001$, and $^{****}p \leq 0.0001$.

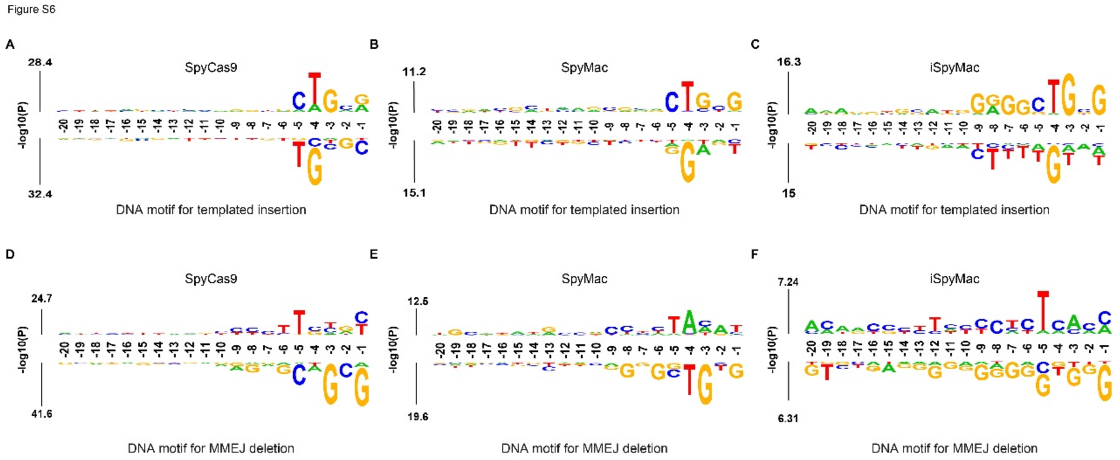


**Figure S6.** Sequence-context motifs associated with templated insertions and MMEJ deletions. (**A-C**) Sequence logos depicting DNA motifs associated with templated insertion for SpyCas9 (A), SpyMac (B), and iSpyMac (C). Enriched nucleotides are shown above the axis and depleted nucleotides below the axis; letter heights indicate statistical significance (−log10 P) from the logo analysis. Positions are numbered relative to PAM, with -20 to -1 corresponding to the 20-nt protospacer sequence upstream of PAM. (**D-F**) Sequence logos depicting DNA motifs associated with MMEJ deletion for SpyCas9 (D), SpyMac (E), and iSpyMac (F), plotted as in (A-C).

Figure S7

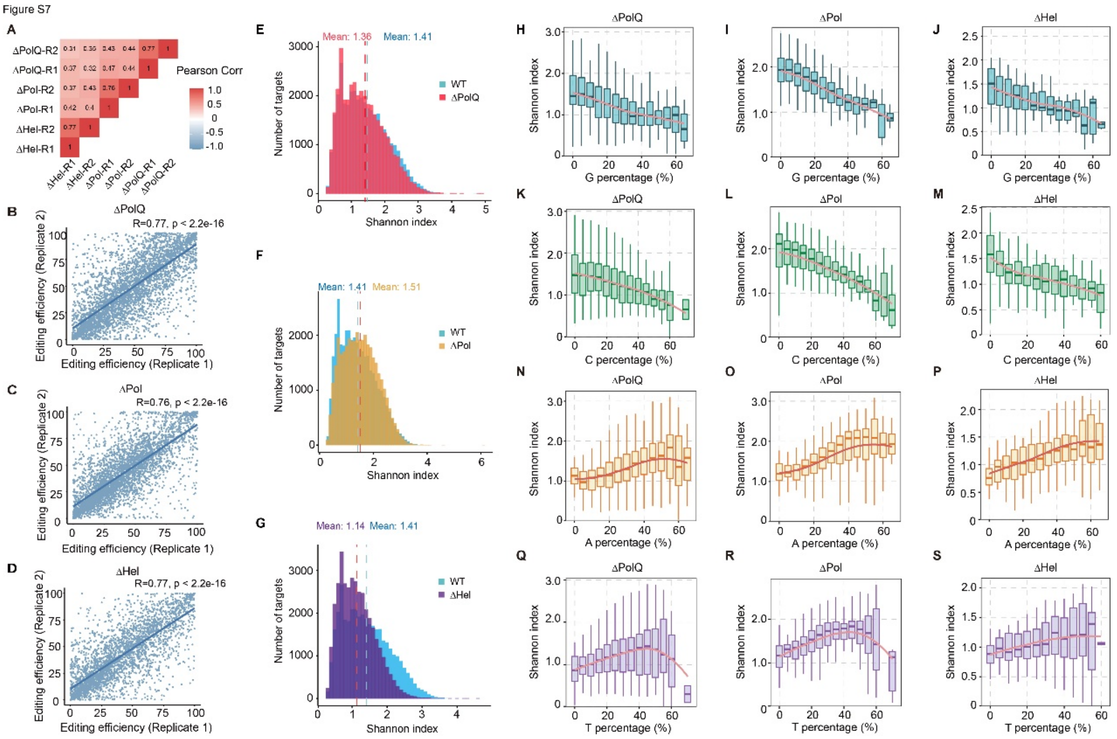


**Figure S7.** Reproducibility and repair-outcome diversity under PolQ deficiency. (**A**) Pearson correlation matrix comparing per-target editing efficiencies across biological replicates under ΔPolQ, ΔPol, and ΔHel conditions. (**B-D**) Replicate concordance of per-target editing efficiencies for ΔPolQ (B), ΔPol (C), and ΔHel (D) with trendlines overlaid; Pearson correlation coefficients and *P* values are shown. (**E-G**) Distributions of repair-outcome diversity across libraries quantified by Shannon index in WT cells compared with ΔPolQ (E), ΔPol (F), and ΔHel (G); dashed lines indicate mean values. (**H-S**) Shannon index as a function of nucleotide composition (G, H-J; C, K-M; A, N-P; T, Q-S) of the 20-nt protospacer under ΔPolQ (H,K,N,Q), ΔPol (I,L,O,R), and ΔHel (J,M,P,S) conditions.

Figure S8

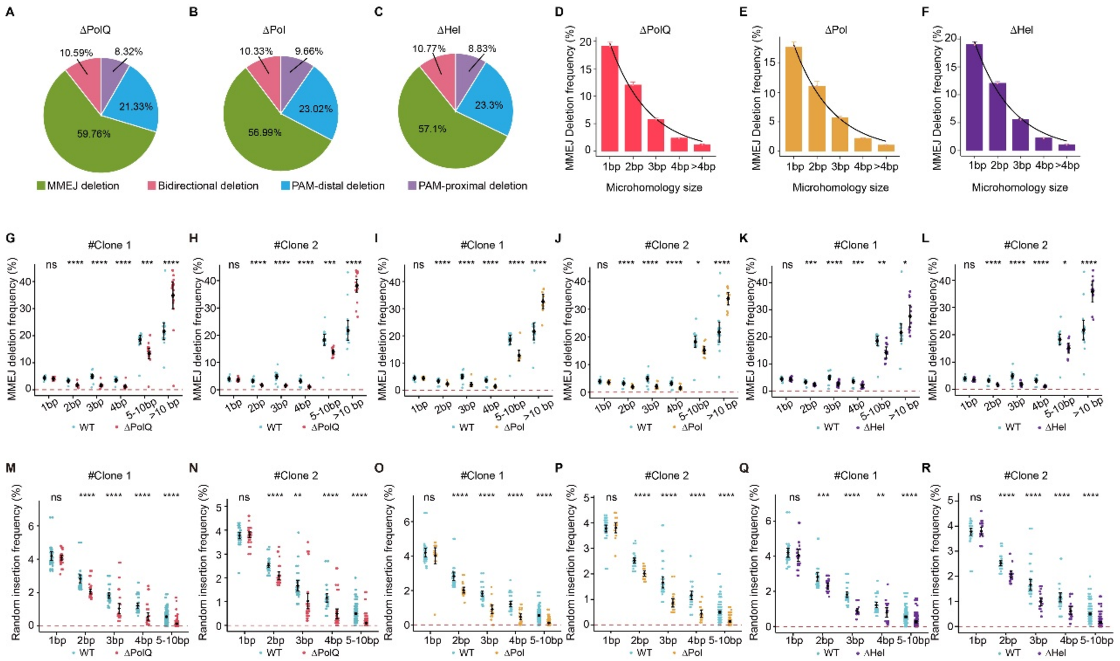


**Figure S8.** Deletion class composition and microhomology dependence under PolQ deficiency. (**A-C**) Deletion profiles of ΔPolQ (A), ΔPol (B), and ΔHel (C). PAM-distal and PAM-proximal deletions were classified relative to the Cas9 cleavage site according to whether the deleted sequence lies on the PAM-distal or PAM-proximal side, respectively. (**D-F**) Microhomology-length usage among MMEJ deletions in ΔPolQ (D), ΔPol (E), and ΔHel (F) cells, shown for 1-4 bp and >4 bp microhomologies with fitted curves. (**G-L**) MMEJ deletion frequencies grouped by deletion size comparing WT with ΔPolQ (G) and (H), ΔPol (**I**) and (J), or ΔHel (K) and (L) for each clone. (**M-R**) Random insertion frequencies grouped by insertion size comparing WT with ΔPolQ (M) and (N), ΔPol (O) and (P), or ΔHel (Q) and (R) for each clone. Statistical significance is indicated and *P* values were calculated in R using an unpaired, two-sided Wilcoxon rank-sum test; **p* ≤ 0.05, ***p* ≤ 0.01, ****p* ≤ 0.001, and *****p* ≤ 0.0001; ns, not significant.

Figure S9

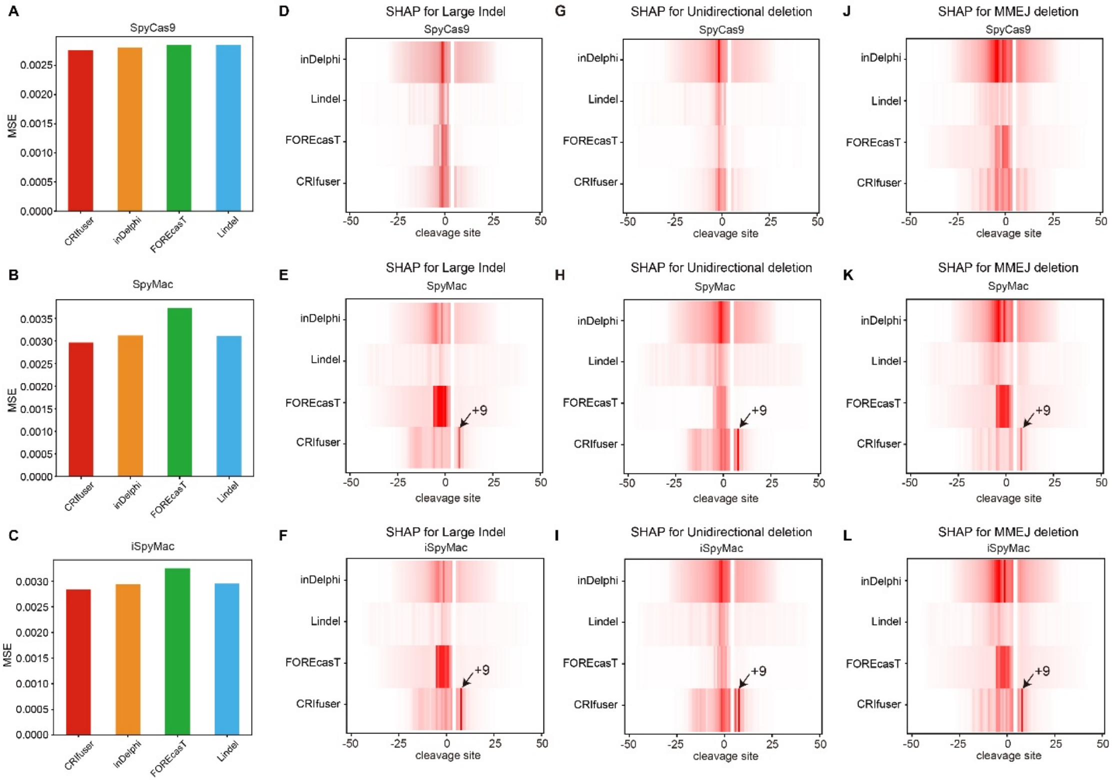


**Figure S9.** CRIfuser predicts distinct SHAP importance across the G-rich and A-rich PAM-recognizing Cas9 editors. (**A-C**) Benchmark for comparison of CRIfuser against inDelphi, FOREcasT, and Lindel measured by mean square error (MSE) for SpyCas9 (A), SpyMac (B), and iSpyMac (C). (**D-L**) SHAP importance of bases around cleavage site in predicting large indel, unidirectional deletion, and MMEJ deletion. Note the strong signals at the +9 position predicted by CRIfuser for the A-rich SpyMac and iSpyMac but not the G-rich SpyCas9. MSE, mean square error.

**Supplementary Note S1: CRIfuser explained.**

**Background of diffusion model.** The diffusion model is widely applied in many disciplines. We develop a diffusion-based model CRIfuser to predict CRISPR/Cas9 editing profiles. Considering that the editing profile is a discrete categorical distribution, we build CRIfuser from the discrete space diffusion model[1,2]. It consists of a U-net architecture and a denoise scheduler for reverse diffusion. We use the target sequence, the cleavage site and the microhomology information as conditional inputs of the diffusion model. Although microhomology information can be inferred from the targeting sequence, we include it directly because it is the dominant mechanism of MMEJ. Start from a randomly selected initial editing outcome, CRIfuser adapt the reverse diffusion of the outcome based on the condition inputs. At each step of the reverse diffusion, the diffusion model predicts the profile of the editing outcome conditioned on the outcome of the previous step. It corrects the profile little by little, and after the whole reverse diffusion, one gets the final predicted editing profile. Actually, the diffusion model is a two-way resemblance of ensemble learning (integrating weak models to obtain a strong model). Firstly, each step in the reverse diffusion uses the output of the previous step to get a better prediction. Secondly, for multiple reverse diffusion processes, the same step predicts the editing profile conditioned on the outcome of the previous step, which is itself sampled from the conditional editing profile of the previous step. That is, the diffusion model performs a weighted bagging repeatedly.

**Microhomology definition.** The editing outcomes are aligned to the cleavage site chimerically. The microhomology happens if the sequence context around the end of the first alignment block and the start of the second alignment block are the same. In this case, there may exist several optimal alignments simultaneously as follows. This is common in MMEJ repair pathway. We shows an example of microhomology of size $3$ below.

```
GACTATAACGCTTGTTAGGATC|GATAGGGATCCGGTACCTTACT   reference

 CTATAACGCTTGTTAGGATCGAT
          CGATAGGGATCCGGTACCTTACT                alignment 1
  CTATAACGCTTGTTAGGATCGA
          TCGATAGGGATCCGGTACCTTACT               alignment 2
   CTATAACGCTTGTTAGGATCG
         ATCGATAGGGATCCGGTACCTTACT               alignment 3
    CTATAACGCTTGTTAGGATC
        GATCGATAGGGATCCGGTACCTTACT               alignment 4
```

**CRIfuser architecture.** The model requires $\pm 25$bp sequence context around the cleavage site. The upstream ligation end $x$ ranges from $-25$bp to $+6$bp. The downstream one $y$ ranges from $-6$bp to $+25$bp. This setting allows the model to predict deletions up to 25bp and templated insertions up to 6bp. The diffusion space of $(x, y)$ is $32 \times 32$. We transform the DNA sequence of $x$ (ranging from $-25$bp to $+6$bp) to the one-hot representation of size $4 \times 31 \times 1$, repeat it along the dimension of $y$ to size $4 \times 31 \times 32$, and pad zeros along the dimension of $x$ to size $4 \times 32 \times 32$. We apply a similar process for the DNA sequence of $y$.The microhomology sizes for all pairs of positions upstream and downstream of the cleavage site are represented by a $25 \times 25$ two-dimensional tensor.We pad it with zeros to size $32 \times 32$ so that it fits the diffusion space of ligation ends. We represent the cleavage site information by a $32 \times 32$ tensor whose elements are all zeros except that of the cleavage site, which takes value $1$. We represent the editing outcome of the previous step by a $32 \times 32$ tensor in the same way as the cleavage site. We then concatenate all the above tensors to a tensor of size $11 \times 32 \times 32$ as the input of the U-net architecture. The down module of the U-net adds the time-step embedding to the input processed by the convolution layer. The softmax layer transforms the output of the U-net model to the editing profile of the current step conditioned on the outcome of the previous step. The full architecture is shown below.

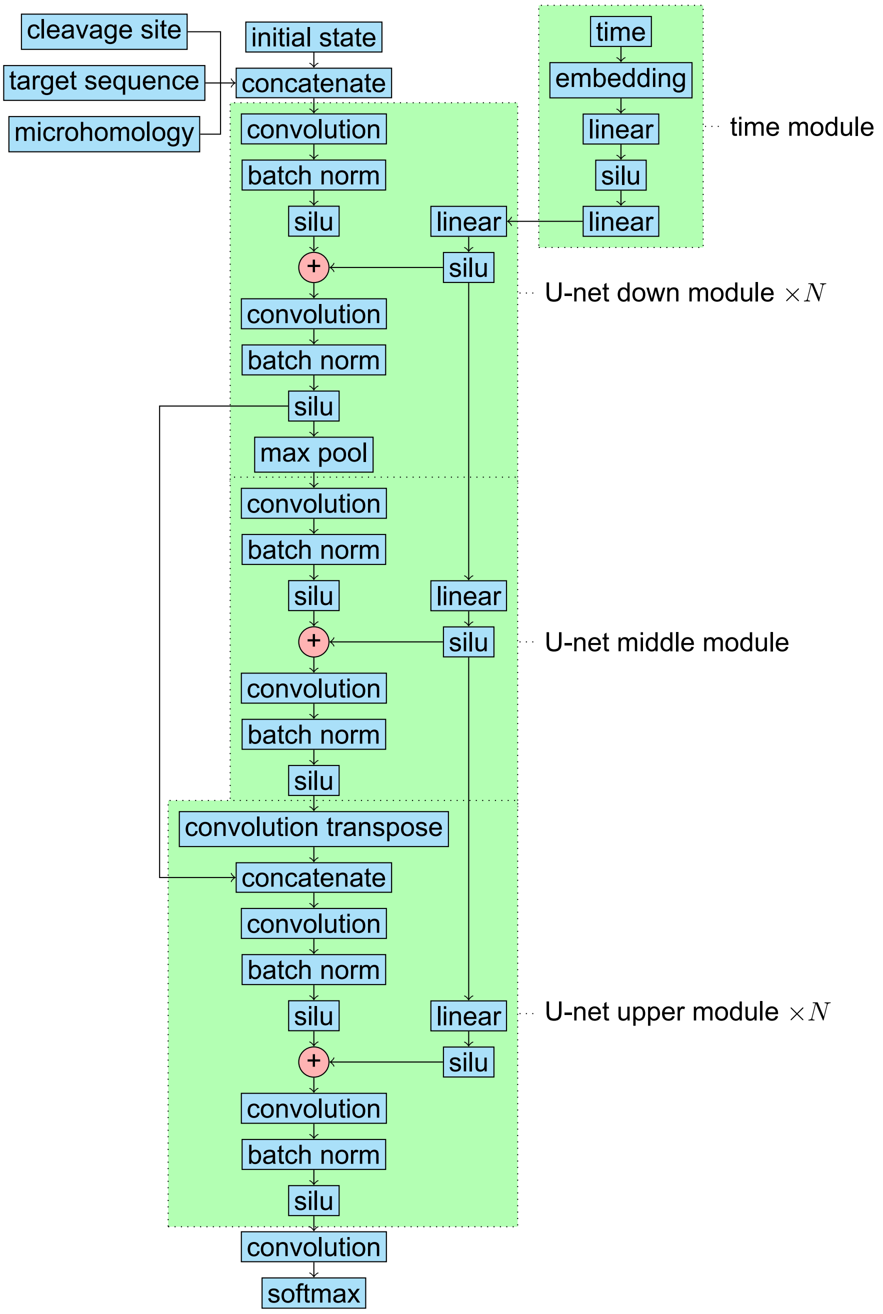


**Prediction of the editing profile.** CRIfuser as well as other diffusion-based models is a sampler by nature. It predicts editing outcomes one at a time. By repeatedly running CRIfuser, one gets lots of editing outcomes which landscape the whole profile. However, the above method is neither efficient nor accurate. To accurately obtain the editing profile, we first predict the editing profile at each step of the reverse diffusion conditioned on all possible outcomes of the previous step, and then sum out the conditioning on the outcome of the previous step by the marginal profile of the previous step. This gives us an accurate marginal profile of the current step. To speed up the prediction of the editing profile, we apply the optimization of contraction order of tensor products[3]. We also implement an interpolation-based approximation of the editing profile. In detail, we perform a grid-sampling with a given step along each dimension of the diffusion space (the upstream and downstream ligation ends). For our $32 \times 32$ diffusion space, a step $n$ will sample $\lfloor 32/n \rfloor \times \lfloor 32/n \rfloor$ outcomes evenly arranged in the diffusion space. Then CRIfuser predicts the editing profile conditioned on all the sampled outcomes. Before summing out the conditioning on the sampled outcomes, we distribute the weights of the non-sampled outcomes to the sampled ones based on the distance-based linear interpolation. We use this interpolation-based approximation exclusively in the Shapley additive explanations.

**Continuous-time discrete-space diffusion model.** We first introduce some notations for the discrete-space diffusion model. We denote the path distribution over continuous time interval $[0, T]$ as $q$. Let $q_t$ be the marginal distribution of $q$ at time step $t$. Let $q_{t|s}$ be the conditional distribution of time step $t$ given time step $s$. The projection of $q$ to a single dimension $1 \leq d \leq D$ is denoted by $q^{(d)}$. The original discrete-space diffusion model is based on discrete time step (DTDS)[4]. Further work generalize the discrete-space diffusion model to continuous time (CTDS)[1,2]. We choose to use the CTDS diffusion model for two reasons.

1. Generally, $q_0$ cannot be decomposed to the product of the marginal distribution of each dimension. Thus, $q_{0|t}$ used in each iteration of the training process of the DTDS diffusion model cannot be decomposed as well. Directly parameterizing $q_{0|t}$ on all dimensions is not efficient, and even infeasible in the high dimensional case for the DTDS diffusion model. The workaround is to parameterize $q_{0|t}(x_0|x_t)$ as the product of marginal distributions $\prod_{d=1}^{D} p_{0|t}^{\theta,(d)}\left(x_0^{(d)}\middle|x_t\right)$, which neglects the dependencies between dimensions. The CTDS diffusion model has not this problem and can parameteriz $q_{0|t}$ smoothly.

2. The DTDS model is trained over specific time steps in a given denoise scheduler. Use a different scheduler from training for inference is possible, but may decrease the accuracy of the model. The CTDS diffusion model is trained over the whole time interval, thereby giving more freedom in the choice of inference denoise scheduler.

Following[2], we use the setting that the diffusion in dimensions are independent, and the forward diffusion process of the diffusion model is

$$q_{t|s}^{(d)}\left(x_t^{(d)}\middle|x_s^{(d)}\right) = \alpha_{t|s}^{(d)}\delta\left(x_t^{(d)}, x_s^{(d)}\right) + \left(1 - \alpha_{t|s}^{(d)}\right)\bar{q}^{(d)}\left(x_t^{(d)}\right), \tag{1}$$

where $\bar{q}^{(d)}$ is the stable distribution of the forward process, and $\lim_{t-s\to+\infty} \alpha_{t|s}^{(d)} = 0$ promises that the forward diffusion process becomes completely random for large time.

This selection greatly simplify the calculation of $q^{(d)}_{s|0,t}\left(x^{(d)}_s \middle| x^{(d)}_0, x^{(d)}_t\right)$[2].

$$
\begin{aligned}
&q^{(d)}_{s|0,t}\left(x^{(d)}_s \middle| x^{(d)}_0, x^{(d)}_t\right) = \frac{q^{(d)}_{t|s}\left(x^{(d)}_t \middle| x^{(d)}_s\right) q^{(d)}_{s|0}\left(x^{(d)}_s \middle| x^{(d)}_0\right)}{q^{(d)}_{t|0}\left(x^{(d)}_t \middle| x^{(d)}_0\right)} = \\
&\begin{cases}
\left(1-\lambda^{(d)}_{t|s}\right)\delta\left(\mathbf{x}^{(d)}_s, \mathbf{x}^{(d)}_t\right) + \lambda^{(d)}_{t|s}\bar{q}^{(d)}\left(x^{(d)}_s\right) & x^{(d)}_0 = x^{(d)}_t \\
\left(1-\mu^{(d)}_{t|s}\right)\delta\left(\mathbf{x}^{(d)}_s, \mathbf{x}^{(d)}_0\right) + \mu^{(d)}_{t|s}\alpha^{(d)}_{t|s}\delta\left(\mathbf{x}^{(d)}_s, \mathbf{x}^{(d)}_t\right) + \mu^{(d)}_{t|s}\left(1-\alpha^{(d)}_{t|s}\right)\bar{q}^{(d)}\left(x^{(d)}_s\right) & x^{(d)}_0 \neq x^{(d)}_t
\end{cases}
\end{aligned} \tag{2}
$$

where

$$
\begin{aligned}
\lambda^{(d)}_{t|s} &:= \frac{\left(1-\alpha^{(d)}_{s|0}\right)\left(1-\alpha^{(d)}_{t|s}\right)\bar{q}^{(d)}\left(x^{(d)}_t\right)}{\alpha^{(d)}_{t|0} + \left(1-\alpha^{(d)}_{t|0}\right)\bar{q}^{(d)}\left(x^{(d)}_t\right)}, \\
\mu^{(d)}_{t|s} &= \frac{1-\alpha^{(d)}_{s|0}}{1-\alpha^{(d)}_{t|0}}.
\end{aligned}
$$

The authors in[1] suggest the Markov forward process ($q^{(d)}_t$ is a row vector)

$$
\frac{dq^{(d)}_t}{dt} = q^{(d)}_t R^{(d)}_t,
$$

where $R^{(d)}_t$ is the transition rate at time $t$. If $R^{(d)}_t$ commutes for all time $t$, then the solution is ($s$ for rows and $t$ for columns)

$$
q^{(d)}_{t|s} = e^{\int_s^t R^{(d)}_\tau d\tau}.
$$

To achieve the commutation condition, they further suggest the forward transition rate $R^{(d)}_t$ of the form (for simplicity, we assume the time-based scale $\beta_t$ is independent of dimension $d$)

$$
R^{(d)}_t := \beta_t \bar{R}^{(d)},
$$

where $\bar{R}^{(d)}$ is a constant transition rate, and $\beta_t$ is a time-dependent scalar. If further require $\left(\bar{R}^{(d)}\right)^2 = \bar{R}^{(d)}$, then

$$
q^{(d)}_{t|s} = e^{\bar{R}^{(d)}\int_s^t \beta_\tau d\tau} = I + (1 - e^{-\int_s^t \beta_\tau d\tau})\bar{R}^{(d)}.
$$

Thus, Eq. (1) is actually

$$
\bar{R}^{(d)} = \mathbf{1}\bar{q}^{(d)} - I, \tag{3}
$$

where $\mathbf{1}$ is a column vector of the size of the total discrete space whose components are all one. To see this, note that

$$
\left(I - \mathbf{1}\bar{q}^{(d)}\right)^2 = I - 2 * \mathbf{1}\bar{q}^{(d)} + \mathbf{1}\bar{q}^{(d)}\mathbf{1}\bar{q}^{(d)} = I - \mathbf{1}\bar{q}^{(d)}.
$$

Therefore,

$$
\begin{aligned}
q^{(d)}_{t|s} &= e^{\bar{R}^{(d)}\int_s^t \beta_\tau d\tau} = I + \left(1 - e^{-\int_s^t \beta_\tau d\tau}\right)\bar{R}^{(d)} \\
&= I + \left(1 - e^{-\int_s^t \beta_\tau d\tau}\right)\left(\mathbf{1}\bar{q}^{(d)} - I\right) = e^{-\int_s^t \beta_\tau d\tau} I + \left(1 - e^{-\int_s^t \beta_\tau d\tau}\right)\mathbf{1}\bar{q}^{(d)},
\end{aligned} \tag{4}
$$

which is Eq. (1) with $\alpha_{t|s}^{(d)} = e^{-\int_s^t \beta_\tau d\tau}$.

According to[2], the transition rate of the reverse diffusion process $\widehat{R}_t$ is calculated by

$$\widehat{R}_t(y_t|x_t) = \sum_{d=1}^{D} R_t^{(d)}\left(x_t^{(d)}\middle|y_t^{(d)}\right)\delta\left(x_t^{(\backslash d)}, y_t^{(\backslash d)}\right) g_t^{(d)}\left(y_t^{(d)}\middle|x_t\right) = \sum_{d=1}^{D} \frac{R_t^{(d)}\left(x_t^{(d)}\middle|y_t^{(d)}\right)\delta\left(x_t^{(\backslash d)}, y_t^{(\backslash d)}\right)}{g_t^{(d)}\left(x_t^{(d)}\middle|y_t\right)}, \tag{5}$$

$$g_t^{(d)}\left(y_t^{(d)}\middle|x_t\right) := \frac{q_t\left(y_t^{(d)}, x_t^{(\backslash d)}\right)}{q_t(x_t)} = \sum_{x_0^{(d)}} \frac{q_{t|0}^{(d)}\left(y_t^{(d)}\middle|x_0^{(d)}\right)}{q_{t|0}^{(d)}\left(x_t^{(d)}\middle|x_0^{(d)}\right)} q_{0|t}^{(d)}\left(x_0^{(d)}\middle|x_t\right), \tag{6}$$

where $x_t^{(\backslash d)}$ means all dimensions except $d$.

**Loss functions overview.** To improve the performance of CRIfuser, we test four negative ELBO loss function to estimate the stepwise local loss of CRIfuser.

1. The double-sampling loss function in Eq. (10)[1].
2. The importance-sampling loss function in Eq. (12)[2].
3. The forward loss function proposed by us in Eq. (15).
4. The reverse loss function proposed by us in Eq. (16).

The double-sampling loss function samples both $x_t$ and $y_t$ in each training step. The importance-sampling loss function samples both $x_0$ and $x_t$. Our forward/reverse loss function only samples $x_t$, thereby decrease the uncertainty of the training process compared to previous negative ELBO loss functions.

We also test two cross-entropy loss functions to estimate the global loss of CRIfuser.

1. The cross-entropy loss function in Eq. (17)[1,2].
2. The integrated cross-entropy loss function proposed by us in Eq. (18).

The cross-entropy loss function samples both $x_0$ and $x_t$ in each training step. Our integrated cross-entropy loss function only samples $x_t$, thereby decrease the uncertainty of the training process.

According to the results in Figure S10, our integrated cross-entropy loss function performs the best among all loss functions over all cas9 datasets. The importance-sampling loss function performs the best among all negative ELBO loss functions. Our reverse loss function achieve lower negative log-likelihood value than the double-sampling loss function, but not as lower as the importance-sampling loss function. Our forward loss function failed to converge the model. The reason needs further study.

Based on the above results, we combine our integrated cross-entropy loss function as the global estimation with the previous importance-sampling loss function as the local estimation and take their mean value as the final loss function for the all formal training of CRIfuser. We give more details of each loss function below.

**Negative ELBO** The authors in[2] suggest to use the negative ELBO as part of the loss function. They parameterize the negative ELBO as ($\Delta t \equiv T/K$ and $t_k \equiv k\Delta t$)

$$\mathcal{L}^{\mathsf{CT}}(\theta) := \lim_{K\to+\infty} \sum_{k=0}^{K-1} -\mathbb{E}_{q_{t_{k+1}}\left(x_{t_{k+1}}\right)} \mathbb{E}_{q_{t_k|t_{k+1}}\left(x_{t_k}|x_{t_{k+1}}\right)} \log\left(\delta\left(x_{t_k}, x_{t_{k+1}}\right) + \widehat{R}^{\theta}_{t_{k+1}}\left(x_{t_k}|x_{t_{k+1}}\right)\Delta t + o(\Delta t)\right)$$

$$= T\mathbb{E}_{\mathcal{U}(t;0,T)}\mathbb{E}_{q_t(x_t)}\left[\sum_{y_t\neq x_t} \widehat{R}^{\theta}_t(y_t|x_t) - \sum_{y_t\neq x_t} R_t(y_t|x_t)\log \widehat{R}^{\theta}_t(x_t|y_t)\right] + C,$$

where

$$\widehat{R}^{\theta}_t(y_t|x_t) := \sum_{d=1}^{D} R^{(d)}_t\left(x^{(d)}_t\middle|y^{(d)}_t\right)\delta\left(x^{(\backslash d)}_t, y^{(\backslash d)}_t\right) g^{\theta,(d)}_t\left(y^{(d)}_t\middle|x_t\right), \tag{7}$$

$$g^{\theta,(d)}_t\left(y^{(d)}_t\middle|x_t\right) := \sum_{x^{(d)}_0} \frac{q^{(d)}_{t|0}\left(y^{(d)}_t\middle|x^{(d)}_0\right)}{q^{(d)}_{t|0}\left(x^{(d)}_t\middle|x^{(d)}_0\right)} p^{\theta,(d)}_{0|t}\left(x^{(d)}_0\middle|x_t\right). \tag{8}$$

Let

$$\mathcal{L}^{\mathsf{CT}}_1(\theta) := T\mathbb{E}_{\mathcal{U}(t;0,T)}\mathbb{E}_{q_t(x_t)} \sum_{y_t\neq x_t} \widehat{R}^{\theta}_t(y_t|x_t),$$

$$\mathcal{L}^{\mathsf{CT}}_2(\theta) := -T\mathbb{E}_{\mathcal{U}(t;0,T)}\mathbb{E}_{q_t(x_t)} \sum_{y_t\neq x_t} R_t(y_t|x_t)\log \widehat{R}^{\theta}_t(x_t|y_t).$$

Then

$$\mathcal{L}^{\mathsf{CT}}(\theta) = \mathcal{L}^{\mathsf{CT}}_1(\theta) + \mathcal{L}^{\mathsf{CT}}_2(\theta) + C.$$

By[2], $\mathcal{L}^{\mathsf{CT}}_1(\theta)$ can be efficiently calculated by

$$\sum_{y_t\neq x_t} \widehat{R}^{\theta}_t(y_t|x_t) = \beta_t \sum_{d=1}^{D} \bar{q}^{(d)}\left(x^{(d)}_t\right) \sum_{y^{(d)}_t} g^{\theta,(d)}_t\left(y^{(d)}_t\middle|x_t\right) + C. \tag{9}$$

**The double-sampling loss function.** In each training iteration of the diffusion model, $x_t$ is fixed through sampling $x_0$, $t$ and $x_t$ sequentially. However, $y_t$ can be any value not equaling $x_t$. The calculation of $\widehat{R}^{\theta}_t(x_t|y_t)$ for each $y_t \neq x_t$ needs an expensive network inference to get $p^{\theta,(d)}_{0|t}(\cdot|y_t)$. The authors of[1] solve this problem by sampling $y_t$ given $x_t$ according to

$$S_t(y_t|x_t) := \frac{R_t(y_t|x_t)}{-R_t(x_t|x_t)}.$$

We call this the double-sampling loss function.

$$\dot{\mathcal{L}}^{\mathsf{CT}}_2(\theta) := T\mathbb{E}_{\mathcal{U}(t;0,T)}\mathbb{E}_{q_t(x_t)}\mathbb{E}_{S_t(y_t|x_t)} R_t(x_t|x_t)\log \widehat{R}^{\theta}_t(x_t|y_t).$$

Note that the double-sampling loss function still needs two forward passes to calculate $\widehat{R}^{\theta}_t(\cdot|x_t)$ for $\mathcal{L}^{\mathsf{CT}}_1(\theta)$ and $\widehat{R}^{\theta}_t(\cdot|y_t)$ for $\dot{\mathcal{L}}^{\mathsf{CT}}_2(\theta)$. To achieve only one forward pass, the authors of[1] approximate $q_t(x_t)$ by $\sum_{x_t} q_t(x_t)S_t(y_t|x_t)$. Then

$$\mathcal{L}^{\mathsf{CT}}_1(\theta) \approx \dot{\mathcal{L}}^{\mathsf{CT}}_1(\theta) := T\mathbb{E}_{\mathcal{U}(t;0,T)}\mathbb{E}_{q_t(x_t)}\mathbb{E}_{S_t(y_t|x_t)} \sum_{z_t\neq y_t} \widehat{R}^{\theta}_t(z_t|y_t).$$

Together with Eqs. (3,7,9),

$$
\begin{aligned}
\dot{\mathcal{L}}_1^{\mathsf{CT}}(\theta) &= T\mathbb{E}_{\mathcal{U}(t;0,T)}\beta_t\mathbb{E}_{q_t(x_t)}\mathbb{E}_{S_t(y_t|x_t)}\sum_{d=1}^{D}\bar{q}^{(d)}\left(y_t^{(d)}\right)\sum_{z_t^{(d)}} g_t^{\theta,(d)}\left(z_t^{(d)}\middle|y_t\right),\\
\dot{\mathcal{L}}_2^{\mathsf{CT}}(\theta) &= -T\mathbb{E}_{\mathcal{U}(t;0,T)}\beta_t\mathbb{E}_{q_t(x_t)}\mathbb{E}_{S_t(y_t|x_t)}\left(\sum_{d=1}^{D}\sum_{z_t^{(d)}\neq x_t^{(d)}}\bar{q}^{(d)}\left(z_t^{(d)}\right)\right)\\
&\log\left(\sum_{d=1}^{D}\bar{q}^{(d)}\left(y_t^{(d)}\right)\delta\left(y_t^{(\backslash d)},x_t^{(\backslash d)}\right)g_t^{\theta,(d)}\left(x_t^{(d)}\middle|y_t\right)\right)+C.
\end{aligned}
\tag{10}
$$

**The importance-sampling loss function.** The authors of[2] use the importance sampling to handle the second term of $\mathcal{L}^{\mathsf{CT}}(\theta)$. The original derivation is quite complex. We give a simplified version.

$$
\begin{aligned}
&T\mathbb{E}_{\mathcal{U}(t;0,T)}\mathbb{E}_{q_t(x_t)}\sum_{y_t\neq x_t}R_t(y_t|x_t)\log\widehat{R}_t^{\theta}(x_t|y_t)\\
&=T\mathbb{E}_{\mathcal{U}(t;0,T)}\mathbb{E}_{q_0(x_0)}\mathbb{E}_{q_{t|0}(x_t|x_0)}\mathbb{E}_{q_{t|0}(y_t|x_0)}[1-\delta(x_t,y_t)]\frac{1}{q_{t|0}(y_t|x_0)}R_t(y_t|x_t)\log\widehat{R}_t^{\theta}(x_t|y_t)\\
&=T\mathbb{E}_{\mathcal{U}(t;0,T)}\mathbb{E}_{q_0(x_0)}\mathbb{E}_{q_{t|0}(y_t|x_0)}\sum_{x_t\neq y_t}\frac{q_{t|0}(x_t|x_0)}{q_{t|0}(y_t|x_0)}R_t(y_t|x_t)\log\widehat{R}_t^{\theta}(x_t|y_t)\\
&=T\mathbb{E}_{\mathcal{U}(t;0,T)}\mathbb{E}_{q_0(x_0)}\mathbb{E}_{q_{t|0}(y_t|x_0)}\sum_{d=1}^{D}\sum_{x_t^{(d)}\neq y_t^{(d)}}\frac{q_{t|0}^{(d)}\left(x_t^{(d)}\middle|x_0^{(d)}\right)}{q_{t|0}^{(d)}\left(y_t^{(d)}|x_0^{(d)}\right)}R_t^{(d)}\left(y_t^{(d)}\middle|x_t^{(d)}\right)\log R_t^{(d)}\left(y_t^{(d)}\middle|x_t^{(d)}\right)g_t^{\theta,(d)}\left(x_t^{(d)}\middle|y_t\right)\\
&=T\mathbb{E}_{\mathcal{U}(t;0,T)}\beta_t\mathbb{E}_{q_0(x_0)}\mathbb{E}_{q_{t|0}(y_t|x_0)}\sum_{d=1}^{D}\bar{q}^{(d)}\left(y_t^{(d)}\right)\sum_{x_t^{(d)}\neq y_t^{(d)}}\frac{q_{t|0}^{(d)}\left(x_t^{(d)}\middle|x_0^{(d)}\right)}{q_{t|0}^{(d)}\left(y_t^{(d)}|x_0^{(d)}\right)}\log g_t^{\theta,(d)}\left(x_t^{(d)}\middle|y_t\right)+C.
\end{aligned}
$$

Exchange $x_t$ and $y_t$ in the last equality, and use the fact that

$$
\log g_t^{\theta,(d)}\left(x_t^{(d)}\middle|x_t\right)=\log\sum_{x_0^{(d)}}\frac{q_{t|0}^{(d)}\left(x_t^{(d)}\middle|x_0^{(d)}\right)}{q_{t|0}^{(d)}\left(x_t^{(d)}\middle|x_0^{(d)}\right)}p_{0|t}^{\theta,(d)}\left(x_0^{(d)}\middle|x_t\right)\equiv\log\sum_{x_0^{(d)}}p_{0|t}^{\theta,(d)}\left(x_0^{(d)}\middle|x_t\right)\equiv\log 1\equiv 0,
\tag{11}
$$

we have the importance-sampling loss function as

$$
\bar{\mathcal{L}}_2^{\mathsf{CT}}(\theta):=-T\mathbb{E}_{\mathcal{U}(t;0,T)}\beta_t\mathbb{E}_{q_{0,t}(x_0,x_t)}\sum_{d=1}^{D}\bar{q}^{(d)}\left(x_t^{(d)}\right)\sum_{y_t^{(d)}}\frac{q_{t|0}^{(d)}\left(y_t^{(d)}\middle|x_0^{(d)}\right)}{q_{t|0}^{(d)}\left(x_t^{(d)}\middle|x_0^{(d)}\right)}\log g_t^{\theta,(d)}\left(y_t^{(d)}\middle|x_t\right)+C.
\tag{12}
$$

**The forward loss function.** We propose a simpler method as follows. By Eq. (5), define the new parameterization of the reverse transition rate as

$$
\tilde{R}_t^{\theta}(x_t|y_t):=\sum_{d=1}^{D}\frac{R_t^{(d)}\left(y_t^{(d)}\middle|x_t^{(d)}\right)\delta\left(x_t^{(\backslash d)},y_t^{(\backslash d)}\right)}{g_t^{\theta,(d)}\left(y_t^{(d)}\middle|x_t\right)}.
\tag{13}
$$

Replace $\widehat{R}_t^\theta(x_t|y_t)$ by $\tilde{R}_t^\theta(x_t|y_t)$ in $\mathcal{L}_2^{\mathsf{CT}}(\theta)$.

$$\widetilde{\mathcal{L}}_2^{\mathsf{CT}}(\theta) := -T\mathbb{E}_{\mathcal{U}(t;0,T)}\mathbb{E}_{q_t(x_t)} \sum_{y_t \neq x_t} R_t(y_t|x_t) \log \tilde{R}_t^\theta(x_t|y_t) + C. \tag{14}$$

Then all one needs is to calculate $g_t^\theta(y_t|x_t)$, which only needs a single pass for $p_{0|t}^\theta(\cdot|x_t)$. We call this the forward loss function.

We find that the second term upon the forward diffusion in Eq. (4) can be efficiently calculated.

$$\begin{aligned} &-\sum_{y_t \neq x_t} R_t(y_t|x_t) \log \tilde{R}_t^\theta(x_t|y_t) = -\sum_{d=1}^{D} \sum_{y_t^{(d)} \neq x_t^{(d)}} R_t^{(d)}\left(y_t^{(d)}\middle|x_t^{(d)}\right) \log \frac{R_t^{(d)}\left(y_t^{(d)}\middle|x_t^{(d)}\right)}{g_t^{\theta,(d)}\left(y_t^{(d)}\middle|x_t\right)} \\ &= \sum_{d=1}^{D} \sum_{y_t^{(d)} \neq x_t^{(d)}} R_t^{(d)}\left(y_t^{(d)}\middle|x_t^{(d)}\right) \log g_t^{\theta,(d)}\left(y_t^{(d)}\middle|x_t\right) + C = \beta_t \sum_{d=1}^{D} \sum_{y_t^{(d)}} \bar{q}^{(b)}\left(y_t^{(d)}\right) g_t^{\theta,(d)}\left(y_t^{(d)}\middle|x_t\right) + C. \end{aligned}$$

Thus, together with Eq. (11), we have

$$\widetilde{\mathcal{L}}_2^{\mathsf{CT}}(\theta) = T\mathbb{E}_{\mathcal{U}(t;0,T)}\beta_t\mathbb{E}_{q_t(x_t)} \sum_{d=1}^{D} \sum_{y_t^{(d)}} \bar{q}^{(b)}\left(y_t^{(d)}\right) g_t^{\theta,(d)}\left(y_t^{(d)}\middle|x_t\right) + C. \tag{15}$$

**The reverse loss function.** By[1] ($\Delta t \equiv T/K$ and $t_k \equiv k\Delta t$),

$$\mathcal{L}^{DT}(\theta) = -\sum_{k=0}^{K-1} \mathbb{E}_{q_{t_k,t_{k+1}}\left(x_{t_k},x_{t_{k+1}}\right)} \log p_{t_k|t_{k+1}}^\theta\left(x_{t_k}|x_{t_{k+1}}\right).$$

The authors of[1] sample $x_{t_k}$, and expand the expectation over $x_{t_{k+1}}$.

$$\begin{aligned} &\mathbb{E}_{q_{t_k,t_{k+1}}\left(x_{t_k},x_{t_{k+1}}\right)} \log p_{t_k|t_{k+1}}^\theta\left(x_{t_k}|x_{t_{k+1}}\right) \\ &= \mathbb{E}_{q_{t_k}\left(x_{t_k}\right)}\mathbb{E}_{q_{t_{k+1}|t_k}\left(x_{t_{k+1}}|x_{t_k}\right)} \log\left(\delta\left(x_{t_k},x_{t_{k+1}}\right) + \widehat{R}_{t_{k+1}}^\theta\left(x_{t_k}|x_{t_{k+1}}\right)\Delta t + o(\Delta t)\right) \\ &= \mathbb{E}_{q_{t_k}\left(x_{t_k}\right)} \sum_{x_{t_{k+1}}} q_{t_{k+1}|t_k}\left(x_{t_{k+1}}|x_{t_k}\right) \log\left(\delta\left(x_{t_k},x_{t_{k+1}}\right) + \widehat{R}_{t_{k+1}}^\theta\left(x_{t_k}|x_{t_{k+1}}\right)\Delta t + o(\Delta t)\right) \\ &= \mathbb{E}_{q_{t_k}\left(x_{t_k}\right)} \sum_{x_{t_{k+1}}} \left(\delta\left(x_{t_k},x_{t_{k+1}}\right) + R_{t_k}\left(x_{t_{k+1}}|x_{t_k}\right)\Delta t + o(\Delta t)\right) \log\left(\delta\left(x_{t_k},x_{t_{k+1}}\right) + \widehat{R}_{t_{k+1}}^\theta\left(x_{t_k}|x_{t_{k+1}}\right)\Delta t + o(\Delta t)\right) \\ &= \mathbb{E}_{q_{t_k}\left(x_{t_k}\right)}\left(\widehat{R}_{t_{k+1}}^\theta(x_{t_k}|x_{t_k})\Delta t + \sum_{x_{t_{k+1}} \neq x_{t_k}} R_{t_k}\left(x_{t_{k+1}}|x_{t_k}\right)\Delta t \log\left(\widehat{R}_{t_{k+1}}^\theta\left(x_{t_k}|x_{t_{k+1}}\right)\Delta t + o(\Delta t)\right) + o(\Delta t)\right) \\ &= \mathbb{E}_{q_{t_k}\left(x_{t_k}\right)}\left(\widehat{R}_{t_{k+1}}^\theta(x_{t_k}|x_{t_k})\Delta t + \sum_{x_{t_{k+1}} \neq x_{t_k}} R_{t_k}\left(x_{t_{k+1}}|x_{t_k}\right)\Delta t\Big(\log \Delta t + \log \widehat{R}_{t_{k+1}}^\theta\left(x_{t_k}|x_{t_{k+1}}\right) + \log(1+o(1))\Big) + o(\Delta t)\right) \\ &= \Delta t\mathbb{E}_{q_{t_k}\left(x_{t_k}\right)}\left(\widehat{R}_{t_{k+1}}^\theta(x_{t_k}|x_{t_k}) + \sum_{x_{t_{k+1}} \neq x_{t_k}} R_{t_k}\left(x_{t_{k+1}}|x_{t_k}\right) \log \widehat{R}_{t_{k+1}}^\theta\left(x_{t_k}|x_{t_{k+1}}\right)\right) + o(\Delta t) + C. \end{aligned}$$

Thus,

$$
\begin{aligned}
\mathcal{L}^{\mathsf{CT}}(\theta) &= \lim_{\Delta t \to 0} \mathcal{L}^{\mathsf{DT}}(\theta) \\
&= -\lim_{\Delta t \to 0} \sum_{k=0}^{K-1} \left[ \Delta t \mathbb{E}_{q_{t_k}(x_{t_k})} \left( \widehat{R}^{\theta}_{t_{k+1}}(x_{t_k}|x_{t_k}) + \sum_{x_{t_{k+1}} \neq x_{t_k}} R_{t_k}\big(x_{t_{k+1}}|x_{t_k}\big) \log \widehat{R}^{\theta}_{t_{k+1}}\big(x_{t_k}|x_{t_{k+1}}\big) \right) + o(\Delta t) + C \right] \\
&= -\int_0^T \mathbb{E}_{q_t(x_t)} \left( \widehat{R}^{\theta}_t(x_t|x_t) + \sum_{y_t \neq x_t} R_t(y_t|x_t) \log \widehat{R}^{\theta}_t(x_t|y_t) \right) dt + C \\
&= \int_0^T \mathbb{E}_{q_t(x_t)} \left( \sum_{y_t \neq x_t} \widehat{R}^{\theta}_t(y_t|x_t) - \sum_{y_t \neq x_t} R_t(y_t|x_t) \log \widehat{R}^{\theta}_t(x_t|y_t) \right) dt + C \\
&= T \mathbb{E}_{\mathcal{U}(t;0,T)} \mathbb{E}_{q_t(x_t)} \left( \sum_{y_t \neq x_t} \widehat{R}^{\theta}_t(y_t|x_t) - \sum_{y_t \neq x_t} R_t(y_t|x_t) \log \widehat{R}^{\theta}_t(x_t|y_t) \right) + C.
\end{aligned}
$$

On the contrary, we choose to sample $x_{t_{k+1}}$, and expand the expectation over $x_{t_k}$.

$$
\begin{aligned}
&\mathbb{E}_{q_{t_k,t_{k+1}}\left(x_{t_k},x_{t_{k+1}}\right)} \log p^{\theta}_{t_k|t_{k+1}}\big(x_{t_k}|x_{t_{k+1}}\big) \\
&= \mathbb{E}_{q_{t_{k+1}}\left(x_{t_{k+1}}\right)} \mathbb{E}_{q_{t_k|t_{k+1}}\left(x_{t_k}|x_{t_{k+1}}\right)} \log \Big( \delta\big(x_{t_k}, x_{t_{k+1}}\big) + \widehat{R}^{\theta}_{t_{k+1}}\big(x_{t_k}|x_{t_{k+1}}\big) \Delta t + o(\Delta t) \Big) \\
&= \mathbb{E}_{q_{t_{k+1}}\left(x_{t_{k+1}}\right)} \sum_{x_{t_k}} q_{t_k|t_{k+1}}\big(x_{t_k}|x_{t_{k+1}}\big) \log \Big( \delta\big(x_{t_k}, x_{t_{k+1}}\big) + \widehat{R}^{\theta}_{t_{k+1}}\big(x_{t_k}|x_{t_{k+1}}\big) \Delta t + o(\Delta t) \Big) \\
&= \mathbb{E}_{q_{t_{k+1}}\left(x_{t_{k+1}}\right)} \sum_{x_{t_k}} \Big( \delta\big(x_{t_k}, x_{t_{k+1}}\big) + \widehat{R}_{t_{k+1}}\big(x_{t_k}|x_{t_{k+1}}\big) \Delta t + o(\Delta t) \Big) \log \Big( \delta\big(x_{t_k}, x_{t_{k+1}}\big) + \widehat{R}^{\theta}_{t_{k+1}}\big(x_{t_k}|x_{t_{k+1}}\big) \Delta t + o(\Delta t) \Big) \\
&= \mathbb{E}_{q_{t_{k+1}}\left(x_{t_{k+1}}\right)} \left( \widehat{R}^{\theta}_{t_{k+1}}\big(x_{t_{k+1}}|x_{t_{k+1}}\big) \Delta t + \sum_{x_{t_k} \neq x_{t_{k+1}}} \widehat{R}_{t_{k+1}}\big(x_{t_k}|x_{t_{k+1}}\big) \Delta t \log \Big( \widehat{R}^{\theta}_{t_{k+1}}\big(x_{t_k}|x_{t_{k+1}}\big) \Delta t + o(\Delta t) \Big) + o(\Delta t) \right) \\
&= \mathbb{E}_{q_{t_{k+1}}\left(x_{t_{k+1}}\right)} \left( \widehat{R}^{\theta}_{t_{k+1}}\big(x_{t_{k+1}}|x_{t_{k+1}}\big) \Delta t + \sum_{x_{t_k} \neq x_{t_{k+1}}} \widehat{R}_{t_{k+1}}\big(x_{t_k}|x_{t_{k+1}}\big) \Delta t \Big( \log \Delta t + \log \widehat{R}^{\theta}_{t_{k+1}}\big(x_{t_k}|x_{t_{k+1}}\big) + \log(1 + o(1)) \Big) \right) \\
&= \Delta t \mathbb{E}_{q_{t_{k+1}}\left(x_{t_{k+1}}\right)} \left( \widehat{R}^{\theta}_{t_{k+1}}\big(x_{t_{k+1}}|x_{t_{k+1}}\big) + \sum_{x_{t_k} \neq x_{t_{k+1}}} \widehat{R}_{t_{k+1}}\big(x_{t_k}|x_{t_{k+1}}\big) \log \widehat{R}^{\theta}_{t_{k+1}}\big(x_{t_k}|x_{t_{k+1}}\big) \right) + o(\Delta t) + C.
\end{aligned}
$$

Thus,

$$
\begin{aligned}
\widehat{\mathcal{L}}^{\mathsf{CT}}(\theta) &= \lim_{\Delta t\to 0}\mathcal{L}^{\mathsf{DT}}(\theta)\\
&= -\lim_{\Delta t\to 0}\sum_{k=0}^{K-1}\left[\Delta t\,\mathbb{E}_{q_{t_{k+1}}\left(x_{t_{k+1}}\right)}\left(\widehat{R}^{\theta}_{t_{k+1}}\left(x_{t_{k+1}}|x_{t_{k+1}}\right)+\sum_{x_{t_k}\neq x_{t_{k+1}}}\widehat{R}_{t_{k+1}}\left(x_{t_k}|x_{t_{k+1}}\right)\log\widehat{R}^{\theta}_{t_{k+1}}\left(x_{t_k}|x_{t_{k+1}}\right)\right)+o(\Delta t)+C\right]\\
&= -\int_0^T\mathbb{E}_{q_t(x_t)}\left(\widehat{R}^{\theta}_t(x_t|x_t)+\sum_{y_t\neq x_t}\widehat{R}_t(y_t|x_t)\log\widehat{R}^{\theta}_t(y_t|x_t)\right)dt+C\\
&= \int_0^T\mathbb{E}_{q_t(x_t)}\left(\sum_{y_t\neq x_t}\widehat{R}^{\theta}_t(y_t|x_t)-\sum_{y_t\neq x_t}\widehat{R}_t(y_t|x_t)\log\widehat{R}^{\theta}_t(y_t|x_t)\right)dt+C\\
&= T\mathbb{E}_{\mathcal{U}(t;0,T)}\mathbb{E}_{q_t(x_t)}\left(\sum_{y_t\neq x_t}\widehat{R}^{\theta}_t(y_t|x_t)-\sum_{y_t\neq x_t}\widehat{R}_t(y_t|x_t)\log\widehat{R}^{\theta}_t(y_t|x_t)\right)+C\\
&= \mathcal{L}^{\mathsf{CT}}_1(\theta)-T\mathbb{E}_{\mathcal{U}(t;0,T)}\mathbb{E}_{q_t(x_t)}\sum_{y_t\neq x_t}\widehat{R}_t(y_t|x_t)\log\widehat{R}^{\theta}_t(y_t|x_t)+C\\
&=: \mathcal{L}^{\mathsf{CT}}_1(\theta)+\widehat{\mathcal{L}}^{\mathsf{CT}}_2(\theta).
\end{aligned}
$$

We call this the reverse loss function. We find that the second term of the reverse loss function can be efficiently calculated as

$$
\begin{aligned}
&-\sum_{y_t\neq x_t}\widehat{R}_t(y_t|x_t)\log\widehat{R}^{\theta}_t(y_t|x_t)\\
&= -\sum_{d=1}^{D}\sum_{y_t^{(d)}\neq x_t^{(d)}}R_t\left(x_t^{(d)}\middle|y_t^{(d)}\right)g_t^{(d)}\left(y_t^{(d)}\middle|x_t\right)\log\left[R_t\left(x_t^{(d)}\middle|y_t^{(d)}\right)g_t^{\theta,(d)}\left(y_t^{(d)}\middle|x_t\right)\right]\\
&= -\sum_{d=1}^{D}\sum_{y_t^{(d)}\neq x_t^{(d)}}R_t\left(x_t^{(d)}\middle|y_t^{(d)}\right)g_t^{(d)}\left(y_t^{(d)}\middle|x_t\right)\log g_t^{\theta,(d)}\left(y_t^{(d)}\middle|x_t\right)+C\\
&= -\beta_t\sum_{d=1}^{D}\bar{q}^{(d)}\left(x_t^{(d)}\right)\sum_{y_t^{(d)}}g_t^{(d)}\left(y_t^{(d)}\middle|x_t\right)\log g_t^{\theta,(d)}\left(y_t^{(d)}\middle|x_t\right)+C.
\end{aligned}
$$

Thus,

$$
\widehat{\mathcal{L}}^{\mathsf{CT}}_2(\theta) = -T\mathbb{E}_{\mathcal{U}(t;0,T)}\beta_t\mathbb{E}_{q_t(x_t)}\sum_{d=1}^{D}\bar{q}^{(d)}\left(x_t^{(d)}\right)\sum_{y_t^{(d)}}\left[g_t^{(d)}\left(y_t^{(d)}\middle|x_t\right)\log g_t^{\theta,(d)}\left(y_t^{(d)}\middle|x_t\right)\right]+C. \tag{16}
$$

**The integrated cross-entropy loss function.** In[1,2], the authors suggest to include cross-entropy loss between $q_{0|t}(x_0|x_t)$ and $p^{\theta}_{0|t}(x_0|x_t)$. In their high dimensional case, they choose to sample both $x_0$ and $x_t$ (it is reasonable to include time-based scale here).

$$
\mathcal{L}^{\mathsf{CE}}(\theta) := -T\mathbb{E}_{\mathcal{U}(t;0,T)}\beta_t\mathbb{E}_{q_t(x_t)}\mathbb{E}_{q_{0|t}(x_0|x_t)}\log p^{\theta}_{0|t}(x_0|x_t) = -T\mathbb{E}_{\mathcal{U}(t;0,T)}\beta_t\mathbb{E}_{q_{0,t}(x_0,x_t)}\log p^{\theta}_{0|t}(x_0|x_t). \tag{17}
$$

We call this the sampling cross-entropy loss. In our 2D case, by the independent assumption of the dimension-wise diffusion,

$$q_{0,t}(x_0, x_t) = q^{(1)}_{t|0}\left(x^{(1)}_t \middle| x^{(1)}_0\right) q_0(x_0) q^{(2)}_{t|0}\left(x^{(2)}_t \middle| x^{(2)}_0\right).$$

For a fixed $x_t$, $q_{0,t}(\cdot, x_t)$ can be calculated in quadratic complexity, and $q_{0|t}(\cdot|x_t)$ can be obtained by normalizing $q_{0,t}(\cdot, x_t)$. This allows to propose a more efficient non-sampling cross-entropy loss as

$$\widehat{\mathcal{L}}^{\mathsf{CE}}(\theta) := -T\mathbb{E}_{\mathcal{U}(t;0,T)}\beta_t \mathbb{E}_{q_t(x_t)} \sum_{x_0} q_{0|t}(x_0|x_t) \log p^{\theta}_{0|t}(x_0|x_t). \tag{18}$$

**Noise scheduler.** We train CRIfuser by the linear scheduler[2]

$$\begin{aligned} \beta_t &= \frac{1}{T-t}, \\ \alpha^{(d)}_{t|s} &= e^{-\int_s^t \beta_\tau d\tau} = \frac{T-t}{T-s}. \end{aligned}$$

**The training algorithm.** According to the theory of the unified discrete diffusion[2],

$$\begin{aligned} & g^{\theta,(d)}_t\left(y^{(d)}_t \middle| x_t\right) \\ &= \frac{1}{\bar{q}^{(d)}\left(x^{(d)}_t\right)} \left[ \left( 1 - \frac{p^{\theta,(d)}_{0|t}\left(x^{(d)}_t \middle| x_t\right)}{1 + \left(\left(\alpha^{(d)}_{t|0}\right)^{-1} - 1\right)\bar{q}^{(d)}\left(x^{(d)}_t\right)} \right) \bar{q}^{(d)}\left(y^{(d)}_t\right) + \frac{\alpha^{(d)}_{t|0}}{1-\alpha^{(d)}_{t|0}} p^{\theta,(d)}_{0|t}\left(y^{(d)}_t \middle| x_t\right) \right] \\ & \left[1 - \delta\left(x^{(d)}_t, y^{(d)}_t\right)\right] + \delta\left(x^{(d)}_t, y^{(d)}_t\right). \end{aligned} \tag{19}$$

Algorithm S1 show the training process.

**Algorithm S1:** The training algorithm.

**Input:** $\bar{q}$, $q_0$, $\lambda$, $T$.

**while** not converges **do**

  Draw $x_0 \sim q_0$;

  Draw $t \sim \mathcal{U}(t; 0, T-1)$;

  **foreach** $d \in [1, D]$ **do**

    Draw $y_t^{(d)} \sim \bar{q}^{(d)}$;

    Draw $b \sim$ Bernoulli$\left(\alpha_{t|0}^{(d)}\right)$;

    $x_t^{(d)} \leftarrow b x_0^{(d)} + (1-b) y_t^{(d)}$;

  Use network to predict $p_{0|t}^{\theta}(\cdot|x_t)$;

  **foreach** $d \in [1, D]$ **do**

    Calculate $g_t^{\theta,(d)}\left(y_t^{(d)}\middle|x_t\right)$ as in Eq. (19);

  Calculate $\widehat{\mathcal{L}}_t^{\mathsf{CT}}(\theta)$ as in Eq. (16);

  **foreach** $d \in [1, D]$ **do**

    $q_{t|0}^{(d)}\left(x_t^{(d)}\middle|x_0^{(d)}\right) \leftarrow \alpha_{t|0}^{(d)} \delta\left(x_0^{(d)}, x_t^{(d)}\right) + \left(1 - \alpha_{t|0}^{(d)}\right) \bar{q}^{(d)}\left(x_t^{(d)}\right)$;

  $q_{0,t}(x_0, x_t) \leftarrow q_0(x_0) \prod_{d=1}^{D} q_{t|0}^{(d)}\left(x_t^{(d)}\middle|x_0^{(d)}\right)$;

  $q_t(x_t) \leftarrow \sum_{x_0} q_{0,t}(x_0, x_t)$;

  $q_{0|t}(x_0|x_t) \leftarrow q_{0,t}(x_0, x_t)/q_t(x_t)$;

  $\widehat{\mathcal{L}}_t^{\mathsf{CE}}(\theta) \leftarrow -\sum_{x_0} q_{0|t}(x_0|x_t) \log p_{0|t}^{\theta}(x_0|x_t)$;

  $\nabla_\theta\left[\widehat{\mathcal{L}}_t^{\mathsf{CT}}(\theta) + \lambda \widehat{\mathcal{L}}_t^{\mathsf{CE}}(\theta)\right]$;

**The sampling algorithm.** Algorithm S2 gives the sampling process through reverse diffusion. We include the MCMC correction in Algorithm S2 [1,2,5]. By [2], the MCMC correct transition probability out of $x_t$ at time step $t$ for a small increment $\Delta t$ is

$$
\begin{aligned}
&\bar{p}_{t,\Delta t}^{\theta,(d)}\left(y_t^{(d)}\middle|x_t\right) \\
&= \Delta t \beta_t \left[\left(2 - \frac{p_{0|t}^{\theta,(d)}\left(x_t^{(d)}\middle|x_t\right)}{1 + \left[\left(\alpha_{t|0}^{(d)}\right)^{-1} - 1\right] \bar{q}^{(d)}\left(x_t^{(d)}\right)}\right) \bar{q}^{(d)}\left(y_t^{(d)}\right) + \frac{\alpha_{t|0}^{(d)}}{1 - \alpha_{t|0}^{(d)}} p_{0|t}^{\theta,(d)}\left(y_t^{(d)}\middle|x_t\right)\right] \\
&\left[1 - \delta\left(x_t^{(d)}, y_t^{(d)}\right)\right].
\end{aligned}
\tag{20}
$$

**Algorithm S2:** The sampling algorithm.

**Input:** $\bar{q}$. $T$. $\Delta t$. $N$.

Draw $x_T \sim q_T$;

**foreach** $t = T-1$ to $1$ **do**

  Get $p^{\theta}_{0|t}(\cdot|x_t)$ by network;

  Draw $x_0 \sim p^{\theta}_{0|t}(\cdot|x_t)$;

  Calculate $q^{(d)}_{t-1|0,t}\left(x^{(d)}_{t-1}\middle|x^{(d)}_0, x^{(d)}_t\right)$ by Eq. (2);

  Draw $x_{t-1} \sim q_{t-1|0,t}(\cdot|x_0, x_t) = \prod_{d=1}^{D} q^{(d)}_{t-1|0,t}\left(\cdot\middle|x^{(d)}_0, x^{(d)}_t\right)$;

  **if** MCMC correction **then**

    $x_{t-1,0} \leftarrow x_{t-1}$;

    **foreach** $n = 1$ to $N$ **do**

      Get $p^{\theta}_{0|t-1}(\cdot|x_{t-1,n-1})$ by network;

      **foreach** $d = 1$ to $D$ **do**

        Calculate $\bar{p}^{\theta,(d)}_{t-1,\Delta t}\left(y^{(d)}_{t-1}\middle|x_{t-1,n-1}\right)$ by Eq. (20);

        Draw $y^{(d)}_{t-1} \sim \bar{p}^{\theta,(d)}_{t-1,\Delta t}(\cdot|x_{t-1,n-1})$;

        Draw $b \sim \sum_{z^{(d)}_{t-1}} \bar{p}^{\theta,(d)}_{t-1,\Delta t}\left(z^{(d)}_{t-1}\middle|x_{t-1,n-1}\right)$;

        $x^{(d)}_{t-1,n} \leftarrow by^{(d)}_{t-1} + (1-b)x^{(d)}_{t-1,n-1}$;

    $x_{t-1} \leftarrow x_{t-1,N}$;

**CRIformer architecture.** Transformer model is good at extracting high-dimensional relationship of the input tokens[6]. For CRISPR/Cas9 editing, this means the micro-homology interactions in the target site. We use the encoder part of the RoFormer architecture with an output layer to predict the CRISPR editing profile[6]. Single bases of the input targeting site sequence are embedded to high dimension space. These embeddings are passed to tandem attention modules and one feed-forward network. The output embedding vector of the last token of the RoFormer encoder is projected to a $32 \times 32$ logits. The final softmax layer transform the logits to the CRISPR editing profile. The component $(x, y)$ in the output profile is the probability of the outcome that the upstream and downstream ligation ends of DSB are $x$ and $y$, respectively. The full architecture of CRIformer is shown below.

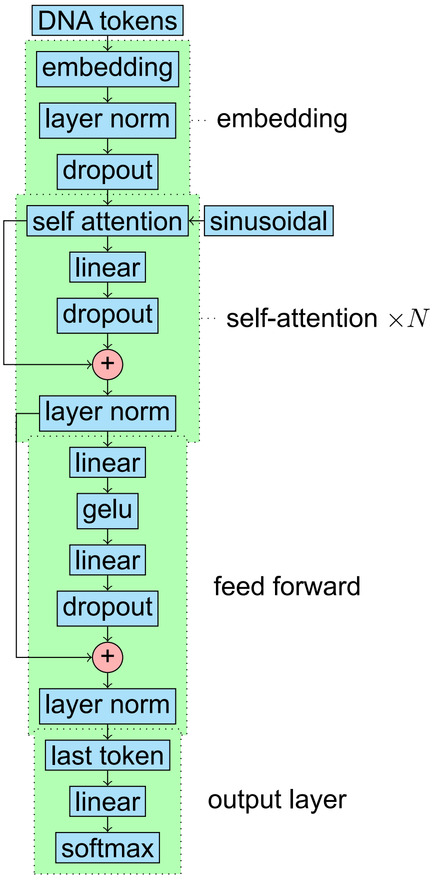


**LSTM architecture.** Resembling[7], we use the 21mer (protospacer + the first nucleotide of PAM) as the input to LSTM model. Besides the 21mer, we also provide some biological inputs to the model, including the free energy of the sgRNA + scaffold, the GC content of the protospacer, the melting temperatures of the whole 21mer input, the first four nucleotides, the middle eight nucleotides from the 5th to the 14th, and the last 5 nucleotides[7]. We embed the 21mer DNA sequence and apply channel-wise dropout. We then feed the result to the bidirectional LSTM layer and activates the output by tanh. We next concatenate the processed sequence input with the biological input and feed them to $N$ (in our case, 3) repeats of linear blocks composed of a linear layer, a elu activation and a dropout layer. Finally, we use an out linear layer to project the embedding space to the $32 \times 32$ logits and use a softmax layer to transform the logits to the predicted editing profile. The full architecture of LSTM is shown below.

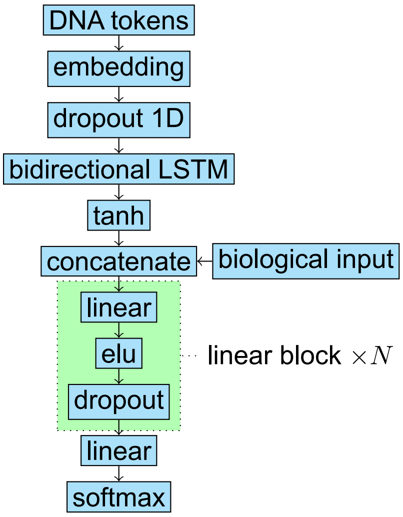


**CNN architecture.** The CNN architecture is similar to the LSTM architecture. The difference is that the bidirectional LSTM becomes parallel CNN blocks. Also, we use the relu activation function to replace the elu in the LSTM architecture (see below).

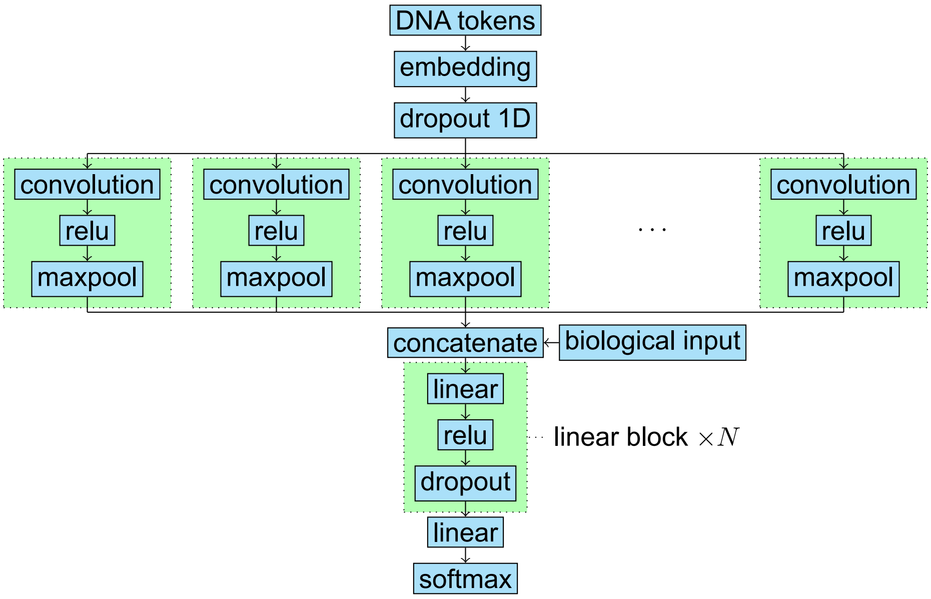


**MLP architecture.** The MLP architecture represents the 21mer DNA as one-hot encoding, and concatenates it with the biological input. Then we simply repeats linear, sigmoid and dropout hidden layers $N$-times (in our setting, 5 times). Finally, we use a linear layer to project the output of hidden layers to the $32 \times 32$ logit tensor and transform it to the editing profile by the softmax layer.

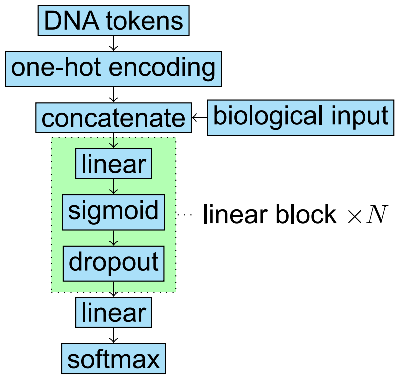
DNA tokens
one-hot encoding
concatenate
biological input
linear
sigmoid
dropout
linear block $\times N$
linear
softmax

**Supplementary Note S2: AI model performances and interpretability.**

**Model architecture selection.** We attempt models of various architectures, including the diffusion model (CRIfuser)[1,2], transformer (CRIformer)[6], long short-term memory (LSTM), convolutional neural network (CNN), multilayer perceptron (MLP), ridge regression and XGBoost, to predict the genome editing profiles including deletion up to 25bp and templated insertion up to 6bp on each side of the cleavage site. We train each model on all three datasets of the G-rich editor SpyCas9, A-rich editor SpyMac and iSpyMac. We find that the diffusion model (CRIfuser) performs the best almost uniformly across all datasets and metrics (Figure S11). The only exception is the mean square error for SpyMac dataset, where CRIfuser performs slightly worse than CRIformer (basically the same).

**Benchmark with models in previous works.** Previous works have tried to predict the CRISPR editing profile from the target sequence, including inDelphi (multilayer perceptron)[8], FOREcasT (logistic regression)[9] and Lindel (logistic regression)[10]. To compare CRIfuser with them, we trained inDelphi, FOREcasT and Lindel on the same datasets as CRIfuser. We conducted the same comparison as in the selection of model architectures. CRIfuser performs uniformly better than models in previous works in all datasets and metrics (Figures 5B-5D and S9A-S9C).

**Shapley additive explanation.** We use SHAP (shapley additive explanation) to explain the model prediction for small indels (indels for both ends of DSB less than or equal to 2bp), large indels (strictly more than 2bp indels for both ends of DSB), unilateral (one end of DSB has an indel less than or equal to 2bp, and the other has one strictly more than 2bp) and MMEJ (large deletions caused by MMEJ). Figure S12 shows the SHAP results for various model architectures. Interestingly, both CRIfuser and CRIformer indicate that the +9 base is very important for the editing outcomes of SpyMac and iSpyMac but not SpyCas9. All other architectures do not give such insights. The models in previous works (inDelphi, FOREcasT, Lindel) do not interpret +9 base as important either (Figures 5E-5G and S9D-S9L).

We also reduce the dimension of SHAP values by UMAP and tSNE Figures S13 and S14. We find that the CRIfuser and CRIformer predict the editing outcomes from the targeting sequence in a similar pattern. On the other hand, traditional machine learning methods like the XGBoost and ridge regression have similar prediction patterns, and their prediction patterns are often quite different from the CRIfuser and CRIformer.

**Supplementary Note S3: The preparation of training data by naapam.**

**Workflow overviews.** The overall workflow to correct the editing profile of CRISPR/Cas9 system is as follows.

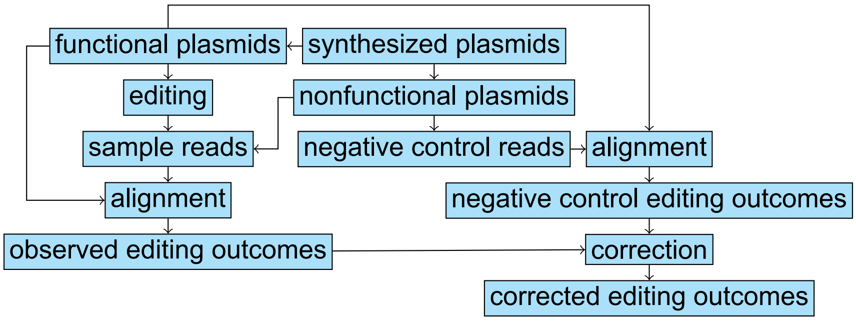


We explain the main steps in details below.

**Parse the sample reads.** The raw sequencing files contain duplicates of the same read. We extract non-duplicated read pairs and reserve the pair counts. Then we parse the elements from the read pairs based on the plasmid design as follows.

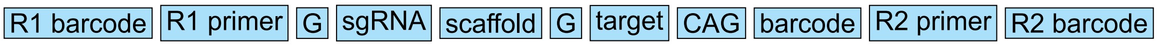


In detail, the plasmid consists of R1 barcode, the last 19 bp of U6 (R1 primer), G, spacer, scaffold, G, 44bp target sequence, CAG, plasmid barcode, 20bp R2 primer and R2 barcode. Read1(R1) sequences the plasmid from U6. Read2(R2) sequences the reverse-complement of the plasmid. Both R1 and R2 are prefixed by sequencing barcode. Because primer and scaffold are fixed across all plasmids, we first extract primer and scaffold sequences from R1 and R2. By using the alignment library of biopython, we allow small mutations in primer and scaffold. For R1, the segment between primer and scaffold is spacer sequence. For R2, that is barcode + CTG + target sequence.

To parse the plasmid barcode in R2, we build bowtie2 index for each plasmid barcode. Then we map the R2 segment between primer and scaffold (*i.e.* barcode + CTG + target sequence) to the plasmid barcodes. Finally, We extract the matching information, including the map score in AS tag, the id of the matched barcode, and the position of the matching.

We use a similar method to parse the spacer in R1 and the protospacer sequence in R2. In detail, we build bowtie2 index for each spacer sequence. Then we map spacer in R1 and target sequence in R2 to the sgRNA sequences. Finally, we extract the matching information, including the map score in AS tag, the id of the matched spacer sequence, and the position of the matching.

**Get the editing targets.** Using the designed plasmids as the editing targets is not accurate because the synthesis of the oligonucleotide is error-prone. Thus, the actual sequence transferred into the cells may be quite different from the design. Instead, we collect the editing targets from the control samples (samples without Cas9 plasmid). We group the control samples by the synthesis libraries, remove duplicates and reserve the counts. These control reads are candidates of editing targets. We filter control reads containing out-of-distribution values of the following criteria.

- Abnormally short or long component length.
- Low alignment score to the designed sequence (e.g. low quality primer, barcode, spacer and scaffold).
- U6 flags necessary for transcription (two Gs padding the sgRNA and scaffold).
- Low count across all control samples.
- Reads without matched barcode/protospacer or with inconsistent barcode and protospacer.

Even after removing duplicates and filtering out low quality control reads, the diversity of the control reads is still too large as editing targets. To further decrease the control read diversity, we align control reads to the designed plasmids to get the synthesis error types of each read, and cluster control reads by synthesis error types. In detail, we neglect irrelevant small synthesis errors away from the cleavage site and only classify control reads by the main synthesis error around the cleavage site, including resections, templated insertions at both ends, and random insertions. We then apply the quality control over the synthesis error types. Firstly, we apply a plasmid barcode-wise filtering. We filter out all synthesis types refer to the designed plasmid of a barcode if the plasmid barcode satisfies any of the following conditions.

- Too many synthesis types.
- With both low total read count and low wild-type frequency.
- The second most common synthesis type is too abundant compared with the most common synthesis type.

Secondly, we filter out synthesis types satisfying any of the following conditions.

- Among all synthesis types of the plasmid of the barcode, the aggregated percentage of those less common than the current synthesis type is low.
- Among all synthesis types of the plasmid of the barcode, the rank of the synthesis type from the most common to the least common is high.
- The upstream or downstream deletion size is too large.
- The random insertion size is too large.
- The count of the synthesis type is too small.

We use the synthesis types passing filtering as editing targets. We align the sequencing reads in treatment samples to these targets instead of the designed sequences to get the editing outcomes.

**The quality control of the sample reads.** We apply a quality control to the sample (both treatment with Cas9 and control without Cas9) reads before aligning them to the editing targets obtained in the previous step. We choose a loose criteria because it is expected that treatment reads will contain more mutants due to the genome editing. In detail, we filter out the sample reads with out-of-distribution values for any of the following criteria.

- Abnormally short or long lengths for R1/R2 primer or scaffold spacer.
- Low alignment score of the designed sequence for R1/R2 primer, scaffold or protospacer.
- The U6 flag necessary for transcription start (the G padding the sgRNA).
- Low count across all samples.
- Reads without matched barcode/protospacer or with inconsistent barcode and protospacer.

**Align the sample reads to the editing targets.** We then assign the sample reads to the editing targets collected from the control samples based on the matched plasmid barcodes. To analyze indel outcomes, especially the templated insertions in high accuracies, we align the sample read to the editing target with the bioconda package rearr (version 1.0.11). If a sample read is assigned to a plasmid barcode with multiple editing targets, we align it to each of the editing target and distribute the count of the sample read to each editing target as follows.

- Normalize the editing target count to get the priory distribution for the count of the sample read.
- Normalize the alignment score between the sample read and the editing target by a temperature and use softmax to calculate the conditional probability.
- By Bayes' theorem, we can get the posterior distribution of the count of the sample read from the priory distribution and the conditional probability.

We collect the alignment results and apply a read-wise filter based on the alignment score. We then aggregate all reads based on the editing outcome near the cleavage site predicted by rearr.

**Filter out the editing target.** During the analyses of editing outcomes, we find that 1bp templated insertion is highly biased to 2bp upstream deletion, 2bp templated insertion is highly biased to 3bp upstream deletion, 3bp templated insertion is highly biased to 4bp upstream deletion and so on. After careful troubleshooting, we find that this is caused by the 1bp deletion synthetic error upstream to the cleavage site. An example of the 3bp templated insertion together with the 4bp upstream deletion caused by the 1bp deletion synthetic error is shown below.

```
CTGGCAAACCACTTATGCTGCTGCT|TCATAAATTTCCAGAATTCTGGCAA
        CTGGCAAACCACTTATGCTGC
                GCTTCATAAATTTCCAGAATTCTGGCAA
```

Because long templated insertion is quite rare, even a low synthesis error rate of 1bp deletion upstream to the cleavage site in the editing target will lead to a great overestimate of the templated insertion. To remove these artifacts, we count reads for each upstream deletion size (not necessarily happening templated insertions) and calculate their relative abundances to the upstream blunt end. We find that the relative abundance decreases exponentially with the upstream deletion size. Thus, we fit a linear model between the logarithm of the relative abundance and the upstream deletion size, as shown in Figure S15.

We calculate the upper bound of the 95% prediction interval as the upper limit of the relative abundance to the upstream blunt end for each upstream deletion size. We expect that these upper limits are good estimates for the relative upstream deletion abundance restricted to any given templated insertion size. We then filter out editing targets with abnormally high abundance of any upstream deletion size restricted to any templated insertion size. We also filter out the editing targets with low total read count or low wild-type frequencies. For the editing targets passing the above criteria, we further filter out their editing outcomes as follows.

- The upstream or downstream deletion size is too large.
- The random insertion size is too large.
- The ratio of the editing target among all editing outcomes is too high.

**Correct the mutant frequency by control sample.** Finally, we apply the correction method used in previous works[11,12]. For each editing target, we calculate its expected abundance in the treatment sample according to its abundance in the control sample. The observed abundance of the editing target will be lower than expected because it is edited by Cas9. The editing outcomes mix with the nonfunctional plasmids. Thus, for each type of the editing outcome, we estimate its expected abundance from the nonfunctional plasmids based on its abundance in the control sample, and subtract the result from the observed abundance. The negative subtraction results are corrected to zero.

**Arrange the data as AI model dataset.** We merge and arrange all alignment results from naapam in a hierarchy JSON format suitable for training process, which is available at https://github.com/ljw20180420/CRISPR_AI/blob/main/AI/dataset/dataset.json.gz. This JSON file is further processed to a Huggingface dataset at https://hf.co/datasets/ljw20180420/CRISPR_AI containing three subsets for each of the SpyCas9, SpyMac and iSpyMac editors. The dataset includes 5'- and 3'-ends of double strand breaks. Each end may undergo resection and templated insertion, thereby shift away from the cleavage site when repair. We train machine learning models to predict the shift of both ends relative to the cleavage site, and reconstruct the genome editing profile based on the shift predictions.

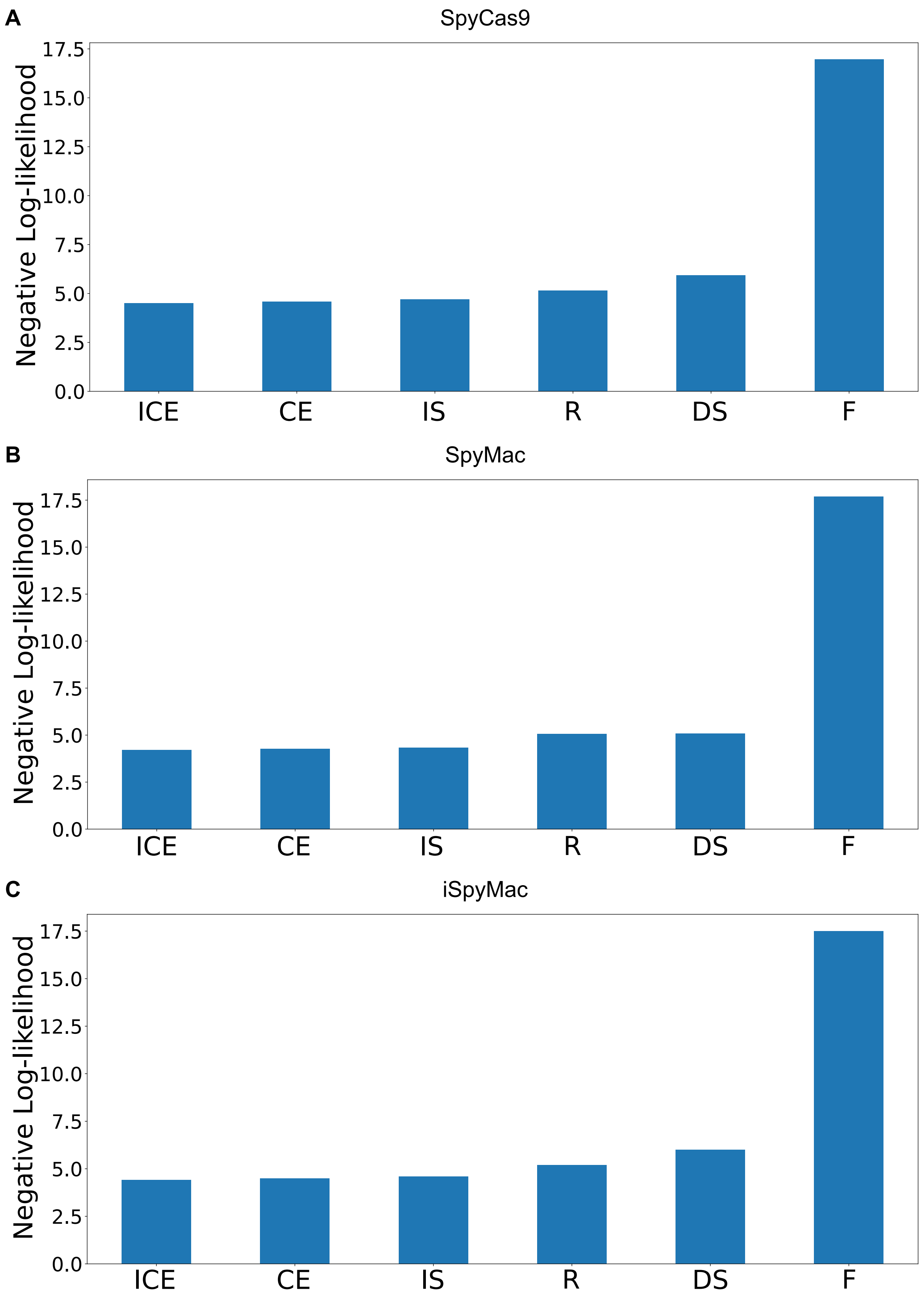


**Figure S10.** Comparing loss functions including the double-sampling loss function (DS), the importance-sampling loss function (IS), the forward loss function (F), the reverse loss function (R), the cross-entropy loss function (CE), and the integrated cross-entropy loss function (ICE). (**A**) SpyCas9 dataset. (**B**) SpyMac dataset. (**C**) iSpyMac dataset.

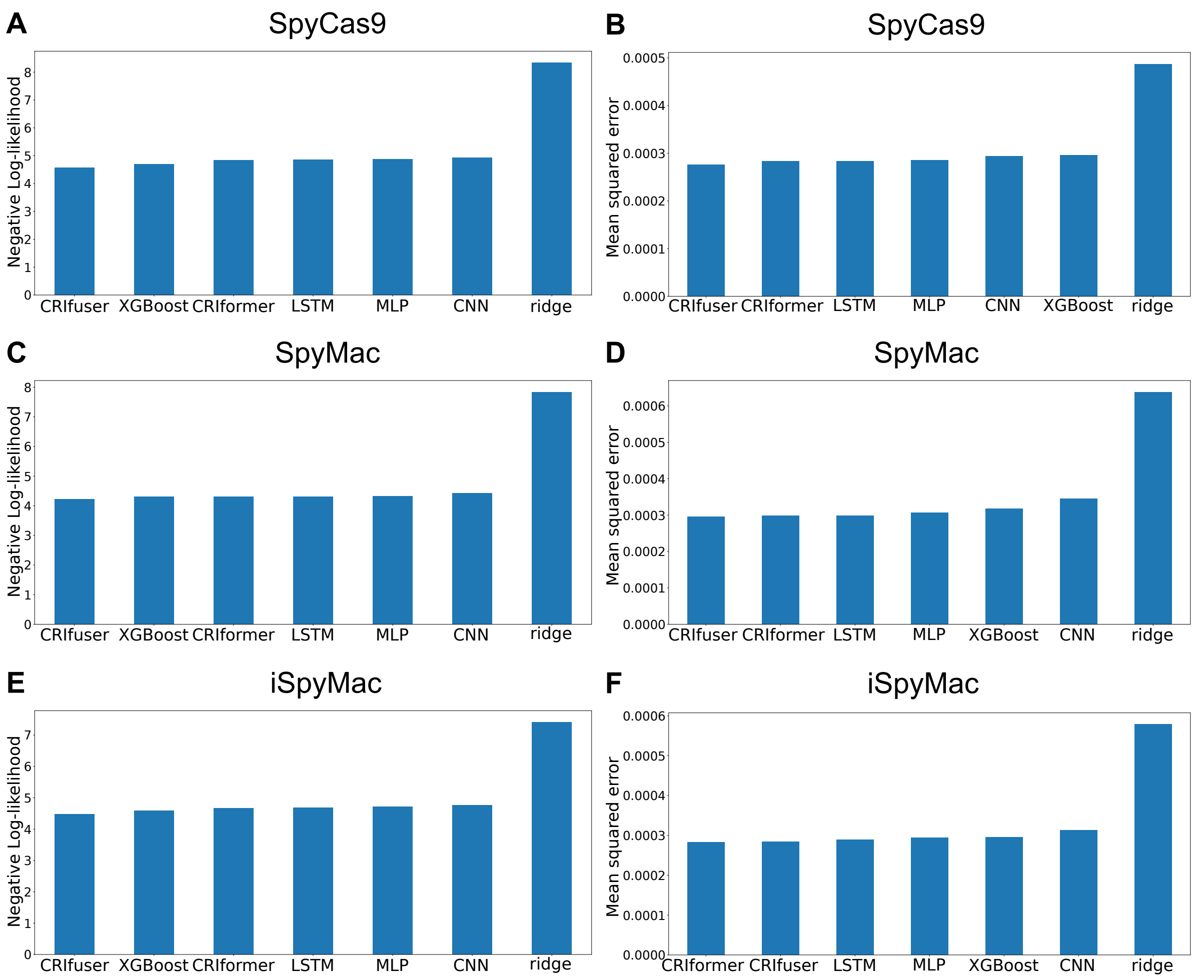


**Figure S11.** Negative log-likelihood and mean squared errors for CRIfuser, XGBoost, CRIformer, LSTM, MLP, CNN and ridge regression on SpyCas9, SpyMac, and iSpyMac datasets. CRIfuser performs the best almost uniformly across all datasets and metrics

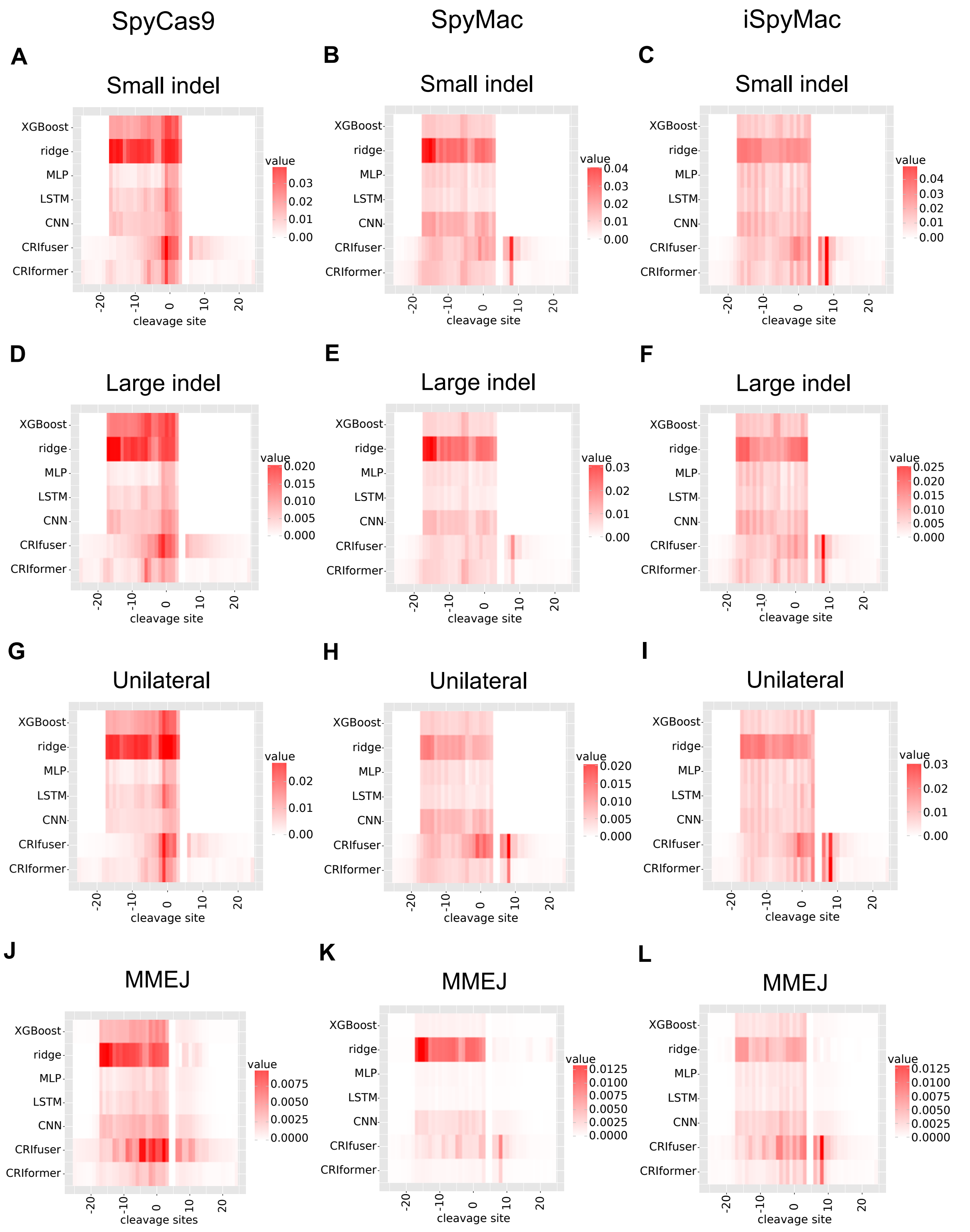


**Figure S12.** Shap results for CRIfuser, CRIformer, LSTM, CNN, MLP, XGBoost and ridge regression on SpyCas9, SpyMac, and iSpyMac datasets. Both CRIfuser and CRIformer predict an important role for +9 position for SpyMac and iSpyMac datasets but not for SpyCas9.

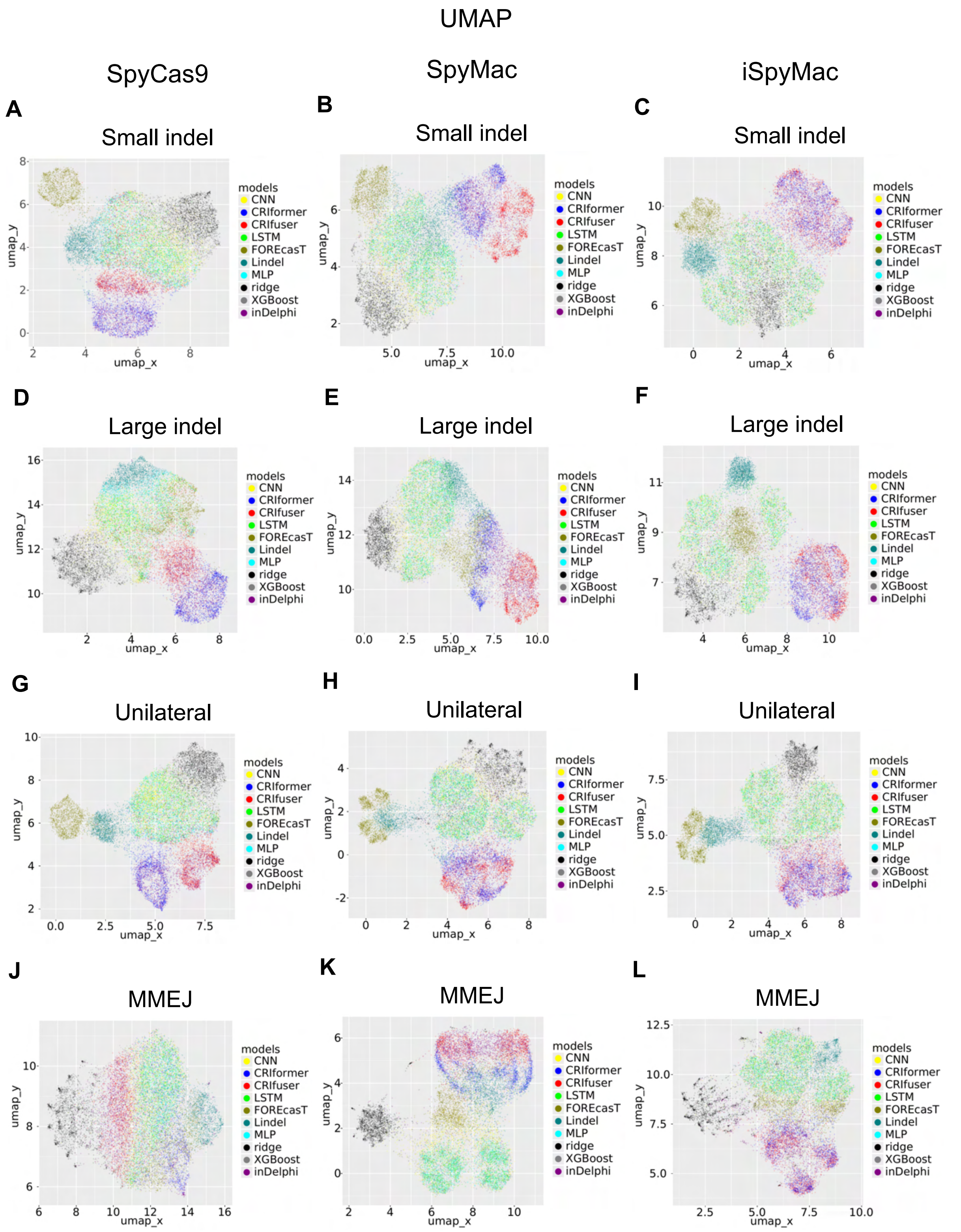


**Figure S13.** SHAP values of different model architectures trained on SpyCas9, SpyMac, and iSpyMac datasets are distinguishable by UMAP. SHAP values of CRIfuser and CRIformer are close to each other and separated from those of the traditional machine learning methods like XG-Boost and ridge regression, except for MMEJ trained on the SpyCas9 dataset.

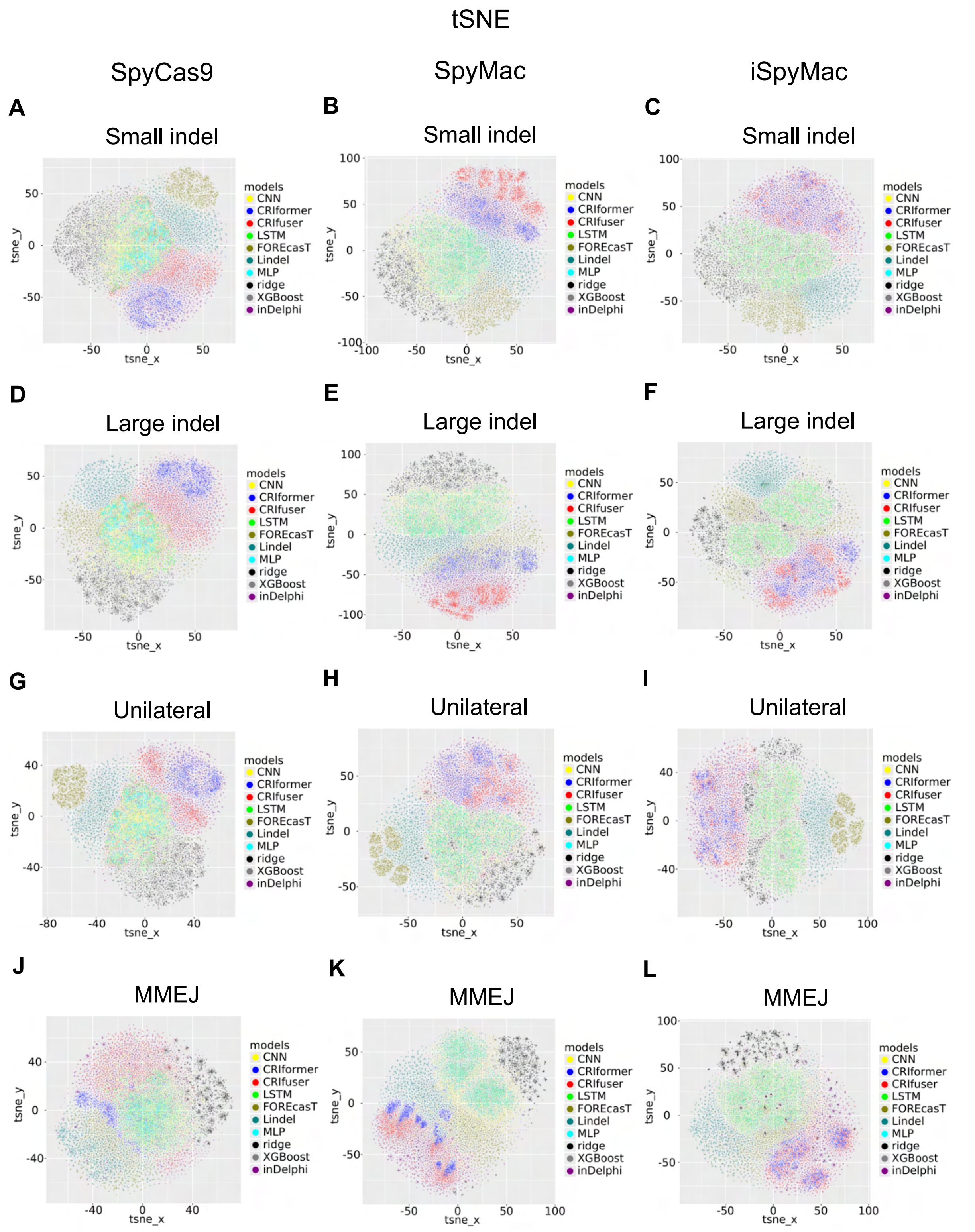


**Figure S14.** SHAP values of different model architectures trained on SpyCas9, SpyMac, and iSpyMac datasets are distinguishable by tSNE. SHAP values of CRIfuser and CRIformer are close to each other and are separated from those of traditional machine learning methods like XG-Boost and ridge regression.

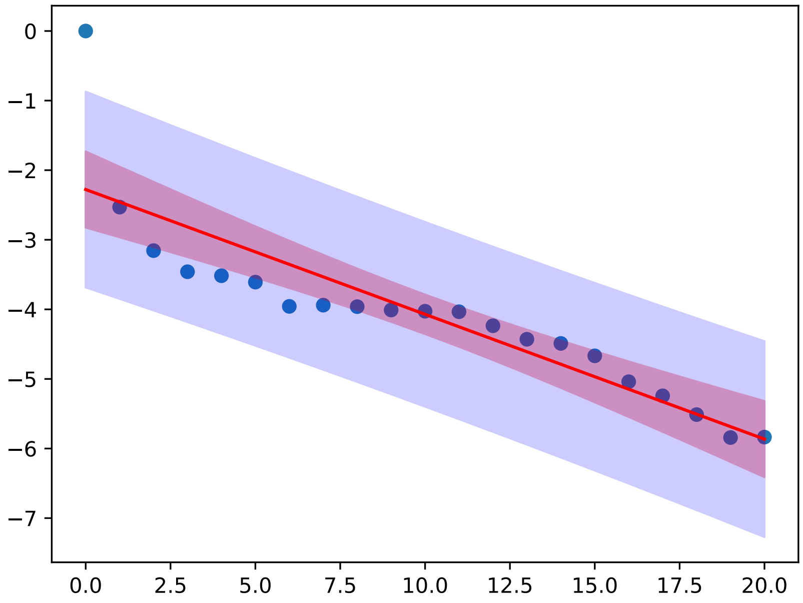


**Figure S15.** The logarithm of the upstream deletion abundance decrease linearly with the deletion size. The deletion abundance is calculated relative to the blunt end.
